# High-resolution transcriptional and morphogenetic profiling of cells from micropatterned human embryonic stem cell gastruloid cultures

**DOI:** 10.1101/2020.01.22.915777

**Authors:** Kyaw Thu Minn, Yuheng C. Fu, Shenghua He, Steven C. George, Mark A. Anastasio, Samantha A. Morris, Lilianna Solnica-Krezel

## Abstract

During mammalian gastrulation, germ layers arise and are shaped into the body plan while extraembryonic layers sustain the embryo. Human embryonic stem cells, cultured with BMP4 on extracellular matrix micro-discs, reproducibly differentiate into gastruloids, expressing markers of germ layers and extraembryonic cells with radial organization. Single-cell RNA sequencing and cross-species comparisons with mouse and cynomolgus monkey gastrulae suggest that gastruloids contain seven major cell types, including epiblast, ectoderm, mesoderm, endoderm, and extraembryonic trophoblast- and amnion-like cells. Upon gastruloid dissociation, single cells reseeded onto micro-discs were motile and aggregated with the same but segregated from distinct cell types. Ectodermal cells segregated from endodermal and extraembryonic cells but mixed with mesodermal cells. Our work demonstrates that the gastruloid system supports primate-specific features of embryogenesis, and that gastruloid cells exhibit evolutionarily conserved sorting behaviors. This work generates a resource for transcriptomes of human extraembryonic and embryonic germ layers differentiated in a stereotyped arrangement.

## Introduction

During mammalian embryogenesis, the first major lineage segregation occurs when trophectoderm (TE), hypoblast (primitive endoderm), and pluripotent epiblast arise in the blastocyst (Cockburn and Rossant, 2010). Whereas epiblast is a precursor of all embryonic tissues, the surrounding hypoblast and TE give rise to extraembryonic (ExE) cells that provide signals to instruct embryonic polarity and germ layers formation (Arnold and Robertson, 2009). TE can be further distinguished into polar (adjacent to epiblast) and mural (distal to epiblast). Despite similarities at the pre-implantation blastocyst stage, developmental differences are observed between post-implantation stages of primate and murine development, as revealed by *in vitro* cultured human and cynomolgus monkey embryos (Deglincerti *et al*., 2016; Shahbazi and Zernicka-Goetz, 2018; Ma *et al*., 2019). Whereas implantation is initiated via polar TE in primates, murine embryos implant using mural TE. Upon implantation in human and primate cynomolgus monkey embryos, epiblast cells adjacent to the polar TE form amnion, while those adjacent to hypoblast form a flat epithelium that will give rise to germ layers during gastrulation. In contrast, germ layers arise from the cup-shaped epiblast epithelium in mouse (Shahbazi and Zernicka-Goetz, 2018).

Gastrulation, during which germ layers are formed and shaped into a body plan, is a fundamental and conserved phase during animal embryogenesis (Solnica-Krezel and Sepich, 2012). During mouse and cynomolgus monkey gastrulation, the primitive streak (PS) forms at the posterior epithelial epiblast, and cells in this region undergo epithelial to mesenchymal transition (EMT) to ingress through the PS and migrate to form mesoderm and endoderm. Cells that remain in the epiblast become ectoderm (Williams *et al*., 2012). Around the same time as gastrulation initiation, primordial germ cells (PGCs) are specified in the epiblast of the mouse embryo (Lawson *et al*., 1999), but likely in the amnion in primates (Sasaki *et al*., 2016).

While EMT endows mesendodermal cells with motility, another morphogenetic process known as cell sorting is thought to underlie proper segregation of germ layers in frog and fish gastrulae. Through cell sorting, germ layers achieve and maintain segregated populations, ensuring establishment of tissue boundaries and organized structures (Fagotto, 2014). First demonstrated by Holtfreter and colleagues, cells from dissociated gastrulating amphibian embryos, when re-aggregated *in vitro*, segregate into their distinct germ layers (Townes and Holtfreter, 1955). Subsequent work in *Xenopus*, *Rana pipiens*, and zebrafish gastrulae further supports reaggregation and segregation of germ layer cells (Davis, Phillips and Steinberg, 1997; Ninomiya and Winklbauer, 2008; Klopper *et al*., 2010). Whether cell sorting is a conserved morphogenetic behavior during mammalian gastrulation, specifically in human, remains to be tested.

Due to ethical and experimental limitations, our knowledge of human gastrulation has largely been derived from studies in model organisms, including mouse and more recently, cynomolgus monkey (Nakamura *et al*., 2016; Ma *et al*., 2019; Niu *et al*., 2019). Internationally recognized guidelines limit studies of human embryogenesis *in vitro* to 14 days post fertilization (dpf) or the onset of gastrulation (Pera, 2017), highlighting the need for alternative *in vitro* models.

Murine embryonic stem cells (ESCs), when cultured in various conditions, not only differentiate into the three germ layers but also recapitulate aspects of spatial patterning, axis formation, symmetry breaking, self-organization, and polarization of gene expression (ten Berge *et al*., 2008; Kopper *et al*., 2009; Fuchs *et al*., 2012; van den Brink *et al*., 2014; Turner *et al*., 2017; Beccari *et al*., 2018). Moreover, mouse epiblast-like cells cultured in micropattern differentiate into spatially-defined regions of cells resembling those found in mouse gastrula (Morgani *et al*., 2018). In more recent studies, co-culture of mouse trophoblast stem cells, ESCs, and ExE endoderm in a 3D matrix generated structures that are morphogenetically, transcriptionally, and structurally similar to natural early murine gastrulae (Harrison *et al*., 2017; Sozen *et al*., 2018). These studies reveal the capability of mouse ESCs to self-organize and recapitulate aspects of *in vivo* gastrulation.

Demonstrating that human pluripotent stem cells (hPSC) may be used to model aspects of human gastrulation, Warmflash *et al*. reported that human ESCs (hESCs), cultured in confined micro-discs of extracellular matrix (ECM) and stimulated with BMP4, can reproducibly differentiate into radially-organized cellular rings, which express markers of ectoderm, mesoderm, endoderm, and TE, arranged from the center outwards, respectively (Warmflash *et al*., 2014). Mesendodermal cells in these cultures exhibit features of EMT, reminiscent of PS formation in murine embryos (Warmflash *et al*., 2014; Martyn, Brivanlou and Siggia, 2019). However, the cellular complexity of the gastruloids remains to be determined.

Here we adapted the micropatterned gastruloid culture developed by Warmflash and colleagues (Warmflash *et al*., 2014) and validated that hESCs in micro-discs, upon BMP4 treatment, reproducibly formed radially organized germ layers and ExE-like cells. We employed single cell RNA sequencing (scRNA-seq) to define the gastruloid cell types and their transcriptomes. These analyses revealed seven cell types, including epiblast, ectodermal, two mesodermal, two endodermal, and ExE-like clusters. The ExE-like cluster is transcriptionally similar to TE and amnion. One of presumptive endodermal clusters contains cells with transcriptional profile similar to early cynomolgus monkey PGC and human PGC-like cells (hPGCLCs). Further cross-species comparisons suggest that the gastruloid cultures correspond to early-to-mid gastrula stage, based on high resemblance in cellular composition to E7.25 mouse and 14 dpf cynomolgus monkey gastrulae. Finally, dissociated gastruloid cells displayed motility and cell sorting behaviors when re-seeded on ECM micro-discs: they tended to aggregate with similar cells, but also to segregate from unlike cell types in a mixed cell population. Thus, our work demonstrates that this simple *in vitro* gastruloid system can generate cells with primate-specific transcriptomes and model conserved morphogenetic cell sorting behaviors of human cells. We also reveal the cellular complexity of BMP4-derived human gastruloids and provide a rich resource for the transcriptomes of embryonic and ExE cells relevant to the gastrulation stage.

## Results

### BMP4 differentiates hESCs in micro-discs into radial patterns of germ layers and ExE cells

To study aspects of human gastrulation, we adapted the micropattern differentiation method developed by Warmflash *et al*. (Warmflash *et al*., 2014) (Figures 1A and S1A–C). After treatment with BMP4 for 44h, H1 hESCs cultured on 500 µm-diameter ECM micro-discs revealed a radial gradient of the downstream effector phosphorylated SMAD1, which declined from the edge to the center (Figure 1B), consistent with previous work (Warmflash *et al*., 2014). Cells in the center expressed the SOX2 transcription factor, a pluripotency marker also expressed during ectodermal differentiation (Li *et al*., 2013). Accordingly, SOX2^+^ cells downregulated the pluripotency marker POU5F1. Surrounding the SOX2^+^ cells was a ring of mesoderm marker Brachyury or T-expressing cells, which were, in turn, abutted by a ring of cells expressing the endoderm marker SOX17 (Figure 1C). We detected two subsets of SOX17^+^ cells, one co-expressing primitive endoderm marker GATA6 and the other, definitive endoderm marker FOXA2 (Figure 1D). The outermost ring contained cells expressing the TE markers CDX2 and GATA3 (Deglincerti *et al*., 2016) (Figures 1C and 1E). ZO-1 expression at the apical cell membranes indicated the epithelial character of these CDX2^+^GATA3^+^ cells (Figure 1E), which also expressed markers of proliferative cytotrophoblasts, p63, TEAD4, and EGFR (Horii *et al*., 2016) (Figure S1E). However, only a subset of them co-expressed KRT7, a marker for pan TE lineage in human (Deglincerti *et al*., 2016) (Figure 1E), suggesting the presence of TE and/or other ExE cellular subtypes in the outermost ring. Taken together, BMP4 treatment of H1 hESCs cultured on ECM micro-discs produced microcolonies termed “gastruloids” with three prospective germ layers surrounded by a ring of ExE-like cells (Figure 1F). As with H1 cells, H9 hESCs showed a similar radial differentiation pattern (Figure S1F).

**Figure 1.**
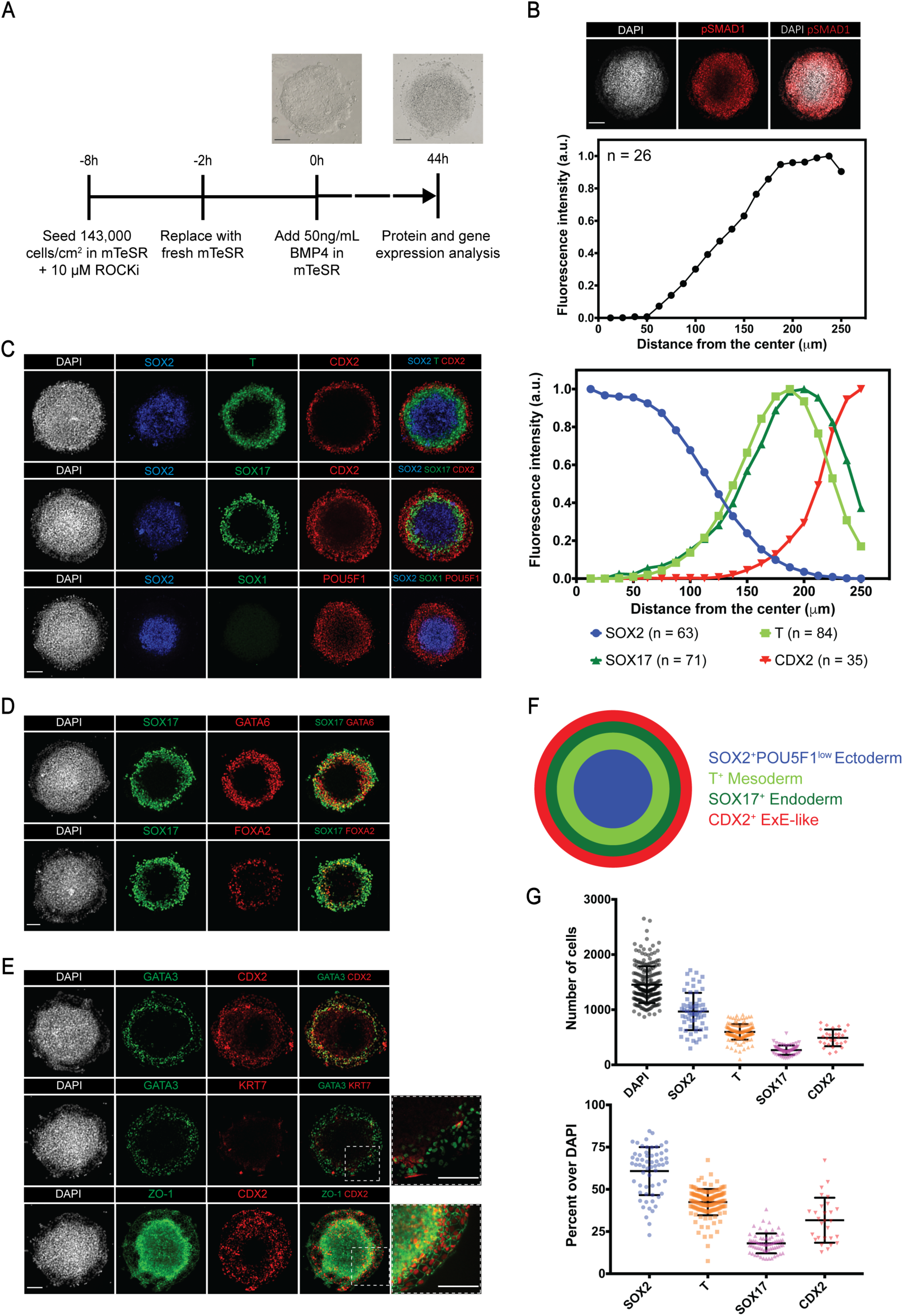
BMP4 induces differentiation of hESCs in Matrigel micro-discs into radially arranged germ layers and ExE cells after 44h. (A) Protocol for BMP4-induced gastruloid differentiation. (B) Immunofluorescence images and quantification of fluorescence intensity for BMP4 downstream effector pSMAD1 relative to the DAPI fluorescence marking nuclei (n = 26). (C) Immunofluorescence images and quantification of fluorescence intensity for indicated markers (n ≥ 35). (D) Immunofluorescence images of primitive and definitive endoderm makers, GATA6 and FOXA2, respectively, in gastruloid cultures. (E) Immunofluorescence images of indicated TE makers in ExE-like population. (F) Schematic representation of cell differentiation in gastruloids. (G) Number of cells expressing indicated markers in gastruloids (n ≥ 28; error bars represent standard deviation). Scale bar is 100 µm.

We assessed the number of cells expressing various markers by using fully convolutional neural network (FCNN) models (see Methods) (He *et al*., 2019). The majority of H1 cells differentiated into SOX2^+^ ectodermal cells (61±14%), followed by T^+^ mesodermal cells (42±8%), CDX2^+^ ExE-like cells (32±13%), and SOX17^+^ endodermal cells (18±6%) (Figure 1G). Both the proportion and the differentiation pattern were consistent across multiple experiments, highlighting the reproducibility of the micropattern culture system (Warmflash *et al*., 2014) (Figures 1C and 1G).

We then asked whether the radial differentiation pattern required BMP4 treatment and micropatterning. We found that BMP4 was sufficient to differentiate germ layer and ExE-like cells in both micropattern and standard monolayer culture (Figures 1C and S1G). However, in the latter, BMP4 treatment resulted in irregularly shaped patterns (Figure S1G). On the other hand, micropattern cultures, without exogenous BMP4, did not generate a pSMAD1 radial gradient (Figure S1D) or express differentiation markers (Figure S1H). Instead, these cells co-expressed SOX2 and POU5F1, indicative of pluripotency (Figure S1H). Taken together, these experiments indicate that BMP4 treatment and micropatterning are required to differentiate hESCs into a stereotyped radial arrangement, consistent with previous reports (Warmflash *et al*., 2014).

### Single-cell profiling reveals additional cell types relevant to gastrulation

Our immunostaining and cell counting data indicated that the total number of cells expressing ectoderm, mesoderm, endoderm, and ExE TE markers exceeded the number of nuclei stained with DAPI (total cells per gastruloid; Figure 1G), suggesting that some cells co-expressed a combination of markers. Accordingly, we detected subsets of cells expressing combinations of endodermal markers (Figure 1D) or TE markers (Figure 1E). Thus, we reasoned that there were more than four distinct cell types and transitional states in the gastruloids.

In order to define all the gastruloid cell types and determine their comprehensive transcriptomes, we performed scRNA-seq on H1 hESC gastruloids after 44h of BMP4 treatment. From two independent experiments, each containing ∼36 gastruloids, we analyzed 2,475 cells with a mean expression of 4,491 genes (Figure S2A). Using the Seurat R toolkit (Butler *et al*., 2018; Stuart *et al*., 2019), we integrated datasets from the two replicates (Figure S2B), resulting in the identification of seven major clusters (Figures 2A and S2C). Using canonical markers, we identified these clusters as Epiblast-like (EPI-like) (*NANOG*^+^*POU5F1*^+^), Ectoderm (*SOX2*^high^*POU5F1*^low^), Mesoderm-1 and −2 (*T^+^MIXL1^+^*), Endoderm-1 and −2 (*SOX17^+^PRDM1^+^*), and ExE-like (TE markers CDX2^+^GATA3^+^ and amnion marker *TFAP2A*^+^) (Figures 2B, 2C, and S2D).

**Figure 2.**
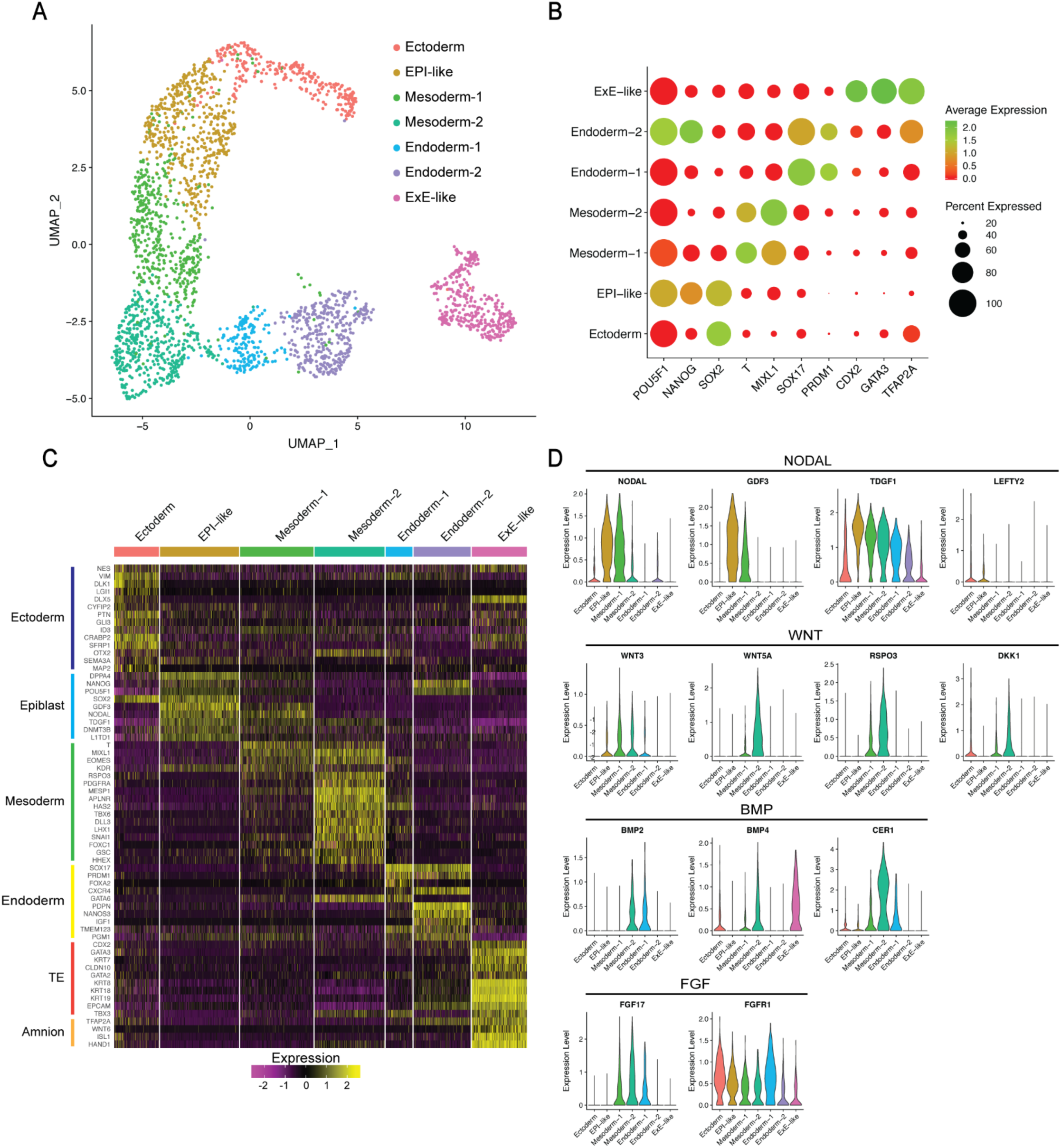
Single-cell RNA-seq profiling reveals cellular complexity of BMP4-induced gastruloids. (A) UMAP display of seven cell clusters detected by scRNA-seq in gastruloid after 44h BMP4 treatment. (B) Dot plot illustrating marker expression of epiblast (*POU5F1, NANOG*), ectoderm (*SOX2*), mesoderm (*T, MIXL1*), endoderm (*SOX17, PRDM1*), TE (*CDX2, GATA3*), and amnion (*TFAP2A*) in the seven gastruloid clusters. (C) Heatmap showing expression of canonical markers of epiblast, ectoderm, mesoderm, endoderm, TE, and amnion in the seven gastruloid clusters. (D) Violin plot showing expression of the FGF, BMP, NODAL, and WNT signaling pathways’ components in the seven gastruloid clusters.

Using Seurat’s single-cell data integration and label transfer functions (Stuart *et al*., 2019), we performed cross-species comparison with scRNA-seq data from gastrulating mouse embryos at E6.5, 6.75, 7.0, 7.25, and E7.5 (Pijuan-Sala *et al*., 2019) and primate cynomolgus monkey at 11, 12, 13, 14, 16, and 17 dpf (Ma *et al*., 2019). Briefly, gastruloid data and mouse or monkey data at all stages were first integrated (Figures 3A–D and S3B–E). Using mouse or monkey data as a reference, their cell type labels were projected onto gastruloid cells, resulting in the labeling of each gastruloid cell with the original cluster annotation (Figure 2A), and predicted mouse or monkey cell types. This allowed us to investigate the composition of predicted mouse or monkey cell types in gastruloids, and whether our originally annotated gastruloid clusters contain relevant mouse or monkey cell types (Figure S3A). In the latter case, we calculated the proportion of predicted mouse or monkey cell types in each originally annotated cluster. We defined a gastruloid cluster as “matching” with certain mouse or monkey cell type if that cluster contained corresponding cell type labels that were computationally predicted. Additionally, permutation tests were used to calculate statistical significance of these cell type “matches” (see Methods).

**Figure 3.**
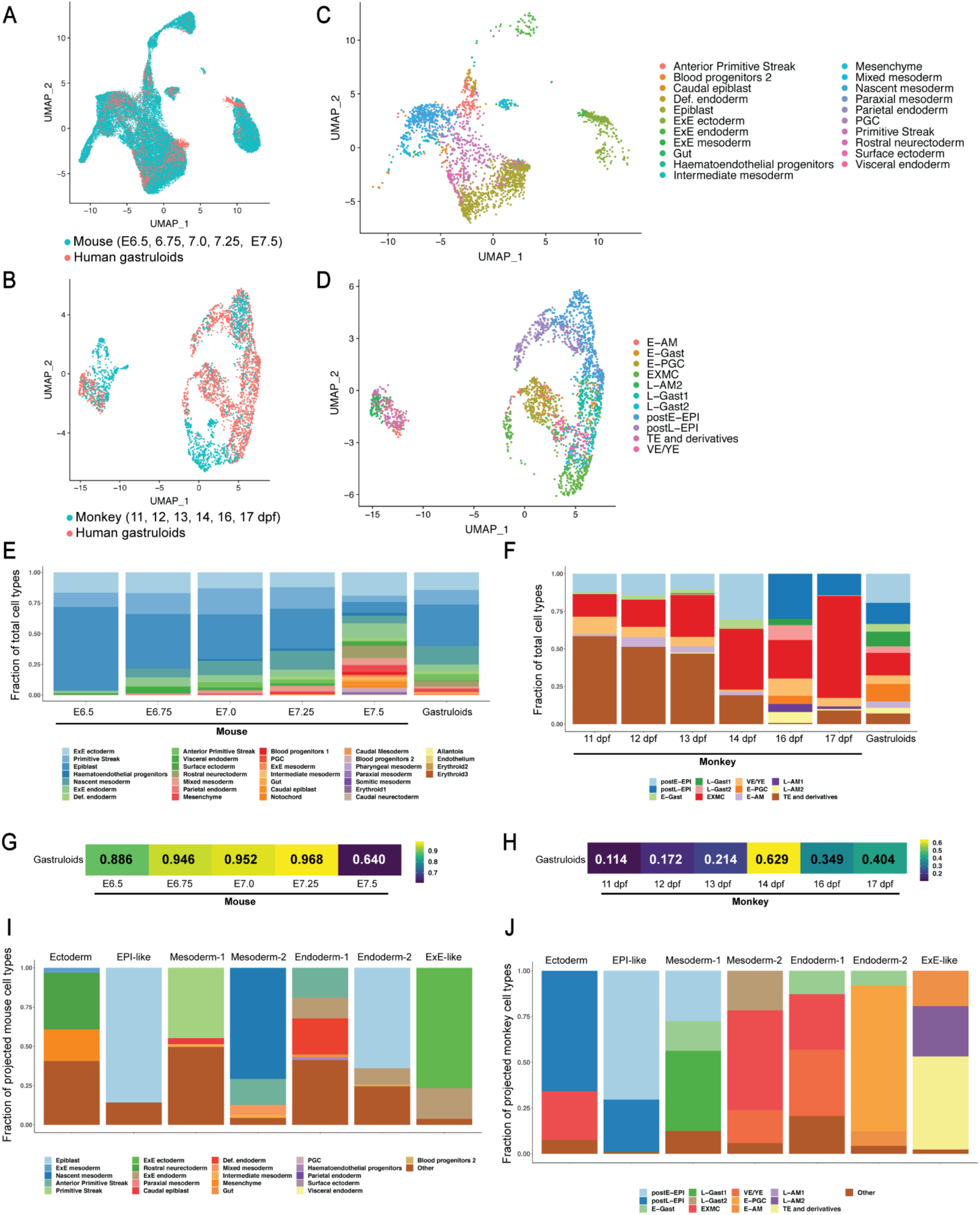
Cross-species comparison with mouse and cynomolgus monkey. (A and B) UMAP displaying overlap of gastruloids and (A) mouse and (B) monkey embryonic cells at indicated stages. (C and D) UMAP showing predicted (C) mouse cell types and (D) monkey cell types in gastruloids. (E and F) Bar plots showing cellular composition of (E) mouse and (F) monkey embryos at indicated stages and gastruloids with predicted cell types. (G and H) Correlation between (G) mouse and (H) monkey developmental stages versus the gastruloid culture based on cell type composition. (I and J) Bar plots showing predicted (I) mouse and (J) monkey cell types in each gastruloid cluster. ‘Other” include cell types that are statistically insignificant using permutation test and *p > 0.05*. Each color bar in E, F, I, and J represents the proportion of a cell type; postE/L-EPI, post-implantation early/late epiblast; E/L-Gast, early/late gastrulating cells; EXMC, ExE mesenchyme; VE/YE, visceral/yolk sac endoderm; E-PGC, early PGC; E/L-AM, early/late amnion.

These analyses revealed that our gastruloid data overlapped with specific mouse and monkey cell types during gastrulation (Figures 3A–D and S3F–G). Comparison of composition between predicted cell types in gastruloids and mouse or monkey cell types at different stages suggested the highest similarity between gastruloids and E7.25 gastrulating mouse (Figures 3E and 3G) or 14 dpf monkey embryo (Figures 3F and 3H), implying that gastruloids likely represent early-to-mid gastrula stage in cellular composition.

### EPI-like cluster with characteristics of post-implantation epiblast

Our scRNA-seq analyses defined a cell cluster enriched for human epiblast markers, *DPPA4*, *NANOG*, *POU5F1*, *SOX2*, *GDF3*, *NODAL*, and *TDGF1* (Petropoulos *et al*., 2016), and mouse epiblast markers, *DNMT3B* and *L1TD1* (Pijuan-Sala *et al*., 2019) (Figure 2C). High activity of NODAL signaling was evidenced by strong expression of *NODAL*, *GDF3*, and *TDGF1* (Figure 2D). We also observed low expression level of *LEFTY2*, a NODAL antagonist and transcriptional target. Since NODAL signaling is known to posteriorize epiblast and is an evolutionarily conserved mesendoderm inducer from fish to mammals (Brennan *et al*., 2001; Schier, 2009), high NODAL activity is consistent with this cluster resembling posterior epiblast. By projecting mouse (Pijuan-Sala *et al*., 2019) or monkey (Ma *et al*., 2019) cell type labels onto the gastruloid data (Stuart *et al*., 2019), we found that the majority of cells in the EPI-like cluster match with mouse epiblast (>80%) (Figure 3I) and monkey post-implantation epiblast (>95%) (Figure 3J). These results suggest that the EPI-like cluster contains cells similar in transcriptomic signature to both monkey and mouse epiblast, and thus precursors of the germ layers.

### Ectodermal cluster expressing nonneural and neural ectoderm markers

Cells in the *SOX2*^high^*POU5F1*^low^ Ectoderm cluster express neuroectoderm markers, *NES* (Lendahl, Zimmerman and McKay, 1990), *VIM* (Schnitzer, Franke and Schachner, 1981), *DLK1* (Surmacz *et al*., 2012), and *LGI1* (Tchieu *et al*., 2017). We also detected transcripts expressed in mouse ectodermal derivatives, *DLX5*, *CYFIP2*, *PTN*, *GLI3*, *ID3*, *CRABP2*, and *SFRP1* (Pijuan-Sala *et al*., 2019) (Figure 2C). While *DLX5*, *CYFIP2, PTN, CRBP2*, and *GLI3* are expressed in rostral neuroectoderm, *DLX5* and *CYFIP2* are also found in surface ectoderm of the E7 mouse embryo. *PTN* and *GLI3* are forebrain/midbrain/hindbrain markers, while *CRBP2* and *SFRP1* are spinal cord markers (Pijuan-Sala *et al*., 2019). Cross-species comparison suggested that cells in this cluster match with mouse rostral neuroectoderm (∼35%) (Figure 3I) and monkey late post-implantation epiblast (∼60%) (Figure 3J), some of which will presumably continue to develop into ectoderm. Interestingly, some cells matched with ExE mesenchyme in mouse (∼20%) and monkey (∼22%). Overall, these data suggest that this cluster resembles ectoderm, expressing nonneural and neural ectoderm genes corresponding to different regional identities along the anterior-posterior axis.

### Mesodermal clusters with characteristics of PS and nascent mesoderm

During gastrulation, mesoderm precursors within the PS, distinguished by the expression of *T* and *MIXL1*, undergo EMT, ingress through the PS, and then migrate to differentiate into paraxial, intermediate, and lateral plate mesoderm (Gilbert, 2010). Our analyses defined two gastruloid mesodermal clusters, termed Mesoderm-1 and Mesoderm-2, both of which expressed markers of PS and mesoderm, *T*, *MIXL1*, and *EOMES* (Costello *et al*., 2011) (Figure 2C). However, Mesoderm-1 expressed *T* at slightly higher levels than Mesoderm-2, suggesting that Mesoderm-1 may represent PS-like cells. Moreover, Mesoderm-2 showed higher transcript levels of genes expressed in mesodermal cells that have already traversed through the PS, including *SNAI1*, and markers of lateral plate *PDGFRA*, *MESP1*, *APLNR*, and *HAS2* (Klewer *et al*., 2006; Vodyanik *et al*., 2010; Chan *et al*., 2013), paraxial *TBX6* and *DLL3* (Lam *et al*., 2014; Loh *et al*., 2016), and intermediate mesoderm *LHX1* (Lam *et al*., 2014) (Figure 2C). Hence, the Mesoderm-2 population likely represents nascent mesoderm and/or precursors of differentiating mesodermal lineages. Consistent with Mesoderm-1 matching mouse PS, cross-species comparison with mouse gastrulae suggested that ∼40% of Mesoderm-1 cells have projected mouse cell type labels of PS. In contrast, Mesoderm-2 matches nascent mesoderm (∼70%) (Figure 3I). Furthermore, cross-species comparison with monkey suggested similarity between Mesoderm-1/2 and monkey gastrulating cells (Figure 3J).

Corroborating previous studies (Warmflash *et al*., 2014; Martyn, Brivanlou and Siggia, 2019), transcriptomes of Mesoderm-1 and −2 contain an EMT signature. A hallmark of EMT during gastrulation is Cadherin switching, where CDH1 is downregulated and CDH2 is upregulated (Hatta and Takeichi, 1986; Radice *et al*., 1997). Immunofluorescence images indicated that T^+^ mesodermal and SOX17^+^ endodermal cells exhibited higher CDH1 in cytoplasm compared to the cell membranes, consistent with CDH1 relocalization from the cell surface to the cytoplasm during EMT (Figure 4A). In contrast, CDH2 immunoreactivity was apparent in T^+^ and SOX17^+^ mesendodermal cells (Figure 4B). Quantification of fluorescence intensity showed that the expression domain of CDH2 largely overlapped with that of T and SOX17 (Figure 4C). As an independent approach, we utilized cosine similarity analysis to measure the spatial overlap in expression patterns between CDH2 and cell type-specific markers (see Methods). Cosine values ranged from 0 to 1, with 0 representing no overlap to 1 representing the highest overlap between expression patterns of two markers. As expected, we found the highest cosine value or overlap between expression patterns of CDH2 and T, followed by that between CDH2 and SOX17 (Figure 4D). In contrast, the cosine values or overlap between expression patterns of CDH2 and GATA3 (ExE-like cells) or SOX2 (ectodermal cells) were significantly lower. These results indicate that T^+^ mesodermal and SOX17^+^ endodermal cells upregulate CDH2, similar to ingressing mesodermal cells in fish, chick, and mouse gastrulae (Hatta and Takeichi, 1986; Radice *et al*., 1997; Warga and Kane, 2007). Consistent with protein expression analyses, we observed high level of *CDH2* but low level of *CDH1* expression in Mesoderm-1/2 and Endoderm-1/2 clusters in scRNA-seq data (Figure 4E).

**Figure 4.**
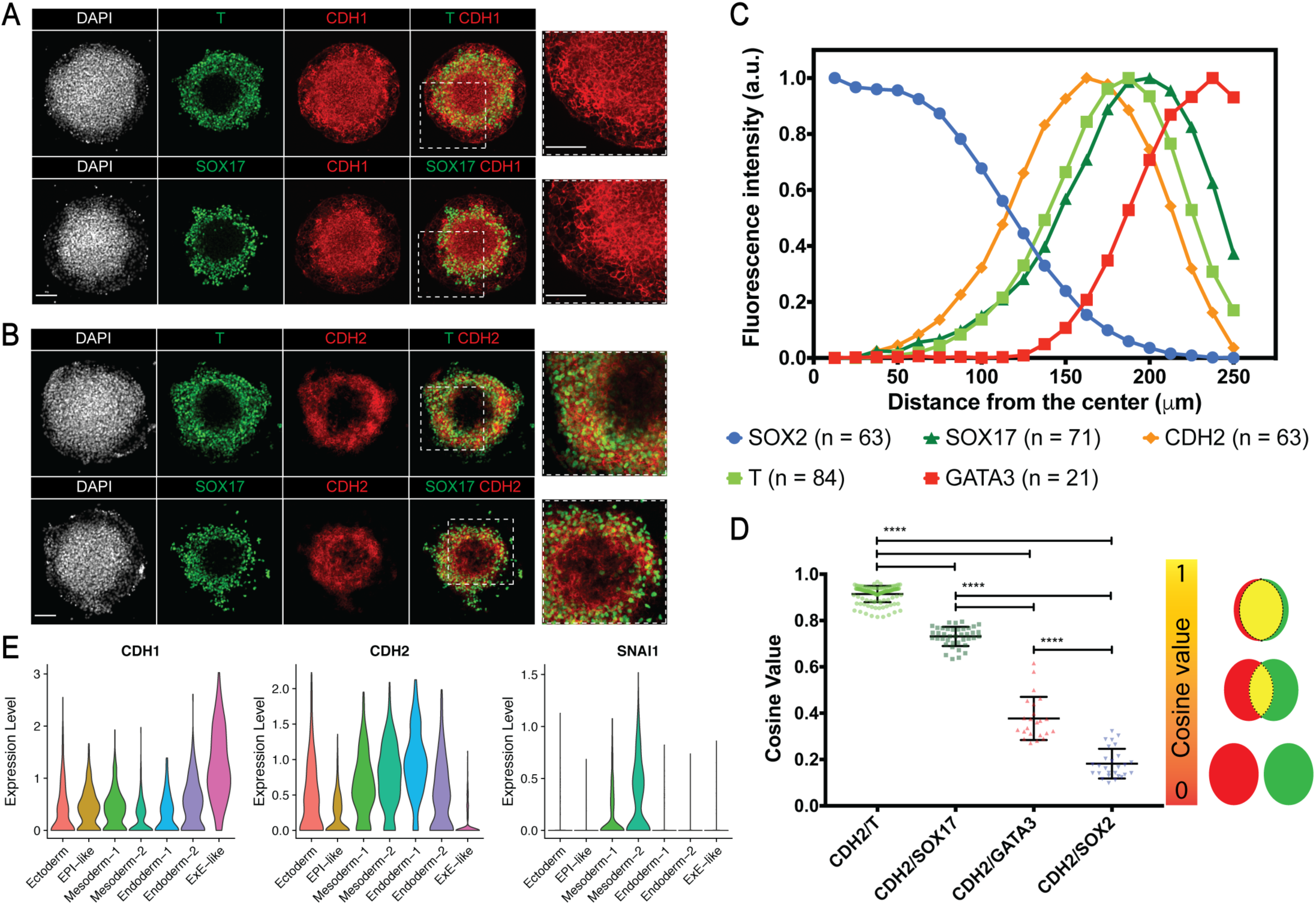
Gastruloid mesendodermal cells show characteristics of cells undergoing EMT in primitive streak. (A and B) Immunofluorescence images of (A) CDH1, and (B) CDH2 in mesendodermal cells (Scale bar 100 μm). (C) Quantification of fluorescence intensity for indicated markers (n ≥ 21). (D) Cosine similarity analysis of expression patterns between CDH2 and indicated marker (n ≥ 23; **** p < 0.0001; Error bars represent standard deviation). (E) Violin plot showing expression of *CDH1*, *CDH2*, and *SNAI1* in the seven gastruloid clusters.

Coincident with CDH2 upregulation in T^+^ mesodermal cells, our scRNA-seq analyses revealed expression of *FGFR1* (Figure 2D) and enrichment of its downstream target *SNAI1* (Warmflash *et al*., 2014) in Mesoderm-1/2 (Figure 4E). SNAI1 is a CDH1 transcriptional repressor and evolutionarily conserved EMT inducer (Barrallo-Gimeno, Nieto and Ip, 2005), suggesting that EMT may be mediated through a similar mechanism in gastruloids. We also found expression of components of FGF (*FGFR1*, *FGF17*), WNT (*WNT3, WNT5A, RSPO3, DKK1*), and TGFβ (*NODAL*, *TDGF1*, *GDF3*) pathways (Figures 2D and S2J), suggesting the activation of these pathways, which has been shown to be responsible for initiation, induction, and maintenance of EMT in mouse (Nieto *et al*., 2016). Interestingly, we observed absent or low level of *FGF8* (not shown), which is dispensable for EMT but required for cell migration away from the PS during mouse gastrulation (Sun *et al*., 1999). Instead, we saw strong expression of *FGF17* in the gastruloid Mesoderm-2 population (Figure 2D), suggesting that a different FGF ligand may be responsible for promoting cell migration during human gastrulation. Hence, these data suggest that while gastruloid cells utilize evolutionarily conserved pathways to undergo EMT and cell migration, specific components involved may differ between human and mouse.

### Gastruloids contain cells resembling primitive and definitive endoderm

We identified two *SOX17*^+^*PRDM1*^+^ presumptive endoderm clusters, termed Endoderm-1 and Endoderm-2 (Figures 2A–C). Endoderm-1 expressed both primitive endoderm marker *GATA6* and definitive endoderm marker *FOXA2* (Figure 2C), consistent with findings from protein expression analysis (Figure 1D). Projection of monkey (Ma *et al*., 2019) cell type labels onto the gastruloid data suggested that Endoderm-1 matches with a mix of visceral endoderm (∼20%), ExE mesenchyme (∼15%), and early/late gastrulating cells (∼20%) (Figure 3J). Similar analysis with mouse showed Endoderm-1 matching with a mix of definitive endoderm (∼20%), anterior PS (∼20%), and ExE endoderm (∼10%) (Figure 3I). Thus, we reasoned that Endoderm-1 cluster represents precursors of both primitive and definitive endoderm.

### Gastruloids contain cells similar in gene expression profiles to primordial germ cells

Cells in Endoderm-2 cluster were enriched for PGC markers *NANOS3* and *TFAP2C* (Sasaki *et al*., 2016; Ma *et al*., 2019; Niu *et al*., 2019; Zheng *et al*., 2019) (Figure S2E), suggesting the presence of hPGCLCs in gastruloids. Additionally, this cluster has a similar gene expression profile to that reported for cynomolgus monkey PGCs (Sasaki *et al*., 2016), expressing *SOX17, TFAP2C, PRDM1, POU5F1*, *NANOG*, and *T*, with low or absent *SOX2* level (Figure S2G–I). The co-expression of SOX17, TFAP2C, NANOG, and NANOS3 as being restricted to hPGCLC has also been reported in a recent study that showed the role of *TFAP2C* in *SOX17* regulation during human germline development (Chen *et al*., 2019). Moreover, cross-species comparison showed Endoderm-2 matching primarily with monkey PGCs (∼80%) (Figure 3J). Interestingly, however, the Endoderm-2 cluster matched primarily with mouse epiblast cells despite showing resemblance to ExE ectoderm and ExE endoderm (Figure 3I). We reasoned that the mismatch occurred because the Endoderm-2 cluster primarily contained hPGCLC, whose formation and gene expression diverge from mouse PGCs (Tan and Tee, 2019). A parallel comparison of our gastruloid data with hPGCLCs derived using *in vitro* human amnion model (Zheng *et al*., 2019), corroborated a strong correlation in gene expression between Endoderm-2 and hPGCLCs in that study (Figure S3H). Together, these findings suggest that the Endoderm-2 cluster contained cells resembling hPGCLCs.

### ExE-like cluster contains cells with gene expression profiles similar to TE and amnion cells

We identified an ExE-like cluster enriched in TE marker genes *CDX2, GATA3*, and *KRT7* (Blakeley *et al*., 2015; Deglincerti *et al*., 2016) (Figure 2C), congruent with findings from protein expression analysis (Figure 1E). The ExE-like cluster also expressed *GATA2* and *TBX3* (Figure 2C), which have been shown to regulate trophoblast differentiation in mouse (Bai *et al*., 2013) and human (Lv *et al*., 2019), respectively. Additionally, *KRT8*/*18*/*19*, found in villous and extravillous trophoblasts of human placenta (Gauster *et al*., 2013), were expressed in the ExE-like cluster. We also noted expression of TE genes observed in E5-7 pre-implantation human embryos (Petropoulos *et al*., 2016), and those in E5 embryos from a separate study (Assou *et al*., 2012) in the ExE-like cluster, including *CLDN4*, *SLC7A2*, and *TACSTD2* (Figure S2F). Therefore, both protein and gene expression analyses suggested the presence of TE-like cells in gastruloids.

Interestingly, the ExE-like cluster also expressed amnion marker genes *TFAP2A*, *HAND1*, *WNT6*, and *ISL1* (Ma *et al*., 2019; Zheng *et al*., 2019) (Figure 2C). Accordingly, comparison with published datasets suggested that cells in ExE-like cluster resemble both TE and amnion cells. Specifically, cross-species comparison with mouse embryo dataset (Pijuan-Sala *et al*., 2019), which do not contain amnion cells, showed cells in ExE-like cluster matching primarily with ExE ectoderm (∼80%) (Figure 3I), a derivative of mouse polar TE (Shahbazi and Zernicka-Goetz, 2018). In contrast, comparison with the *in vitro* human amnion model (Zheng *et al*., 2019), lacking TE cells, showed strong correlation between gastruloid ExE-like cells and amnion-like cells (Figure S3H). Reconciling these results, comparison with monkey embryos, which possess both TE and amnion, showed ExE-like gastruloid cells matching both TE (∼55%) and amnion (∼45%) (Figure 3J). However, further analysis of this ExE-like cluster did not define two distinct populations of TE-like and amnion-like cells (not shown). Taken together, these results suggest that gastruloid differentiation gives rise to ExE-like cells that are transcriptionally similar to both TE and amnion.

### BMP4-differentiated germ layer and ExE-like cells exhibit cell sorting behaviors

During amphibian and fish gastrulation, upon leaving the blastopore (PS equivalent), cells migrate to form distinct layers and organ rudiments. Cell sorting, based on differential cortical tension, adhesion, and/or inhibitory signaling, is thought to be a key driver of these early morphogenetic cell behaviors (Krens and Heisenberg, 2011; Fagotto, 2015; Winklbauer and Parent, 2017). Given the emergence in gastruloids of germ layer and ExE-like cells in variably overlapping and separate radial domains (Figure 1C), we hypothesized that these cells would possess the capability to sort and segregate in a mixed cell population. Therefore, we set out to test whether human gastruloid cells undergo cell sorting *in vitro* as demonstrated in amphibians and fish (Fagotto, 2015).

To this end, H1 gastruloid cultures after 44h BMP4 treatment were dissociated into single cells. The resulting single cell suspension was reseeded onto 500 µm-diameter ECM micro-discs at 72,000 cells/cm^2^ in mTeSR with no additional exogenous factors (Figure 5A). Time-lapse analyses of re-seeded cultures revealed that many cells were actively migrating on the ECM substrate and started to form aggregates by 15h (Figure S4A, Movie S1). Immunostaining at 2h after reseeding revealed a random salt-and-pepper distribution of ectoderm (SOX2^+^), mesoderm (T^+^), endoderm (SOX17^+^), and ExE-like (CDX2^+^) cells (Figure 5B). However, by 24h post-reseeding, cells tended to form homogeneous aggregates with those of the same type. In particular, SOX2^+^, T^+^, SOX17^+^, and CDX2^+^ cells showed a tendency to aggregate with those expressing the same marker. By 48h, such sorting behavior was pronounced for germ layer cells, whereas CDX2^+^ cells tended to surround the germ layer cell clusters. By 72h, while aggregation of SOX2^+^, T^+^, and SOX17^+^ cells was still observable, CDX2^+^ cells mostly dominated individual microcolonies (Figures 5B and S4B–D). As reseeded gastruloid cells preferentially formed aggregates with a similar cell type, we posit that human gastruloid cells display a cell sorting behavior similar to fish and amphibian embryonic cells *in vitro* (Townes and Holtfreter, 1955; Krens and Heisenberg, 2011).

**Figure 5.**
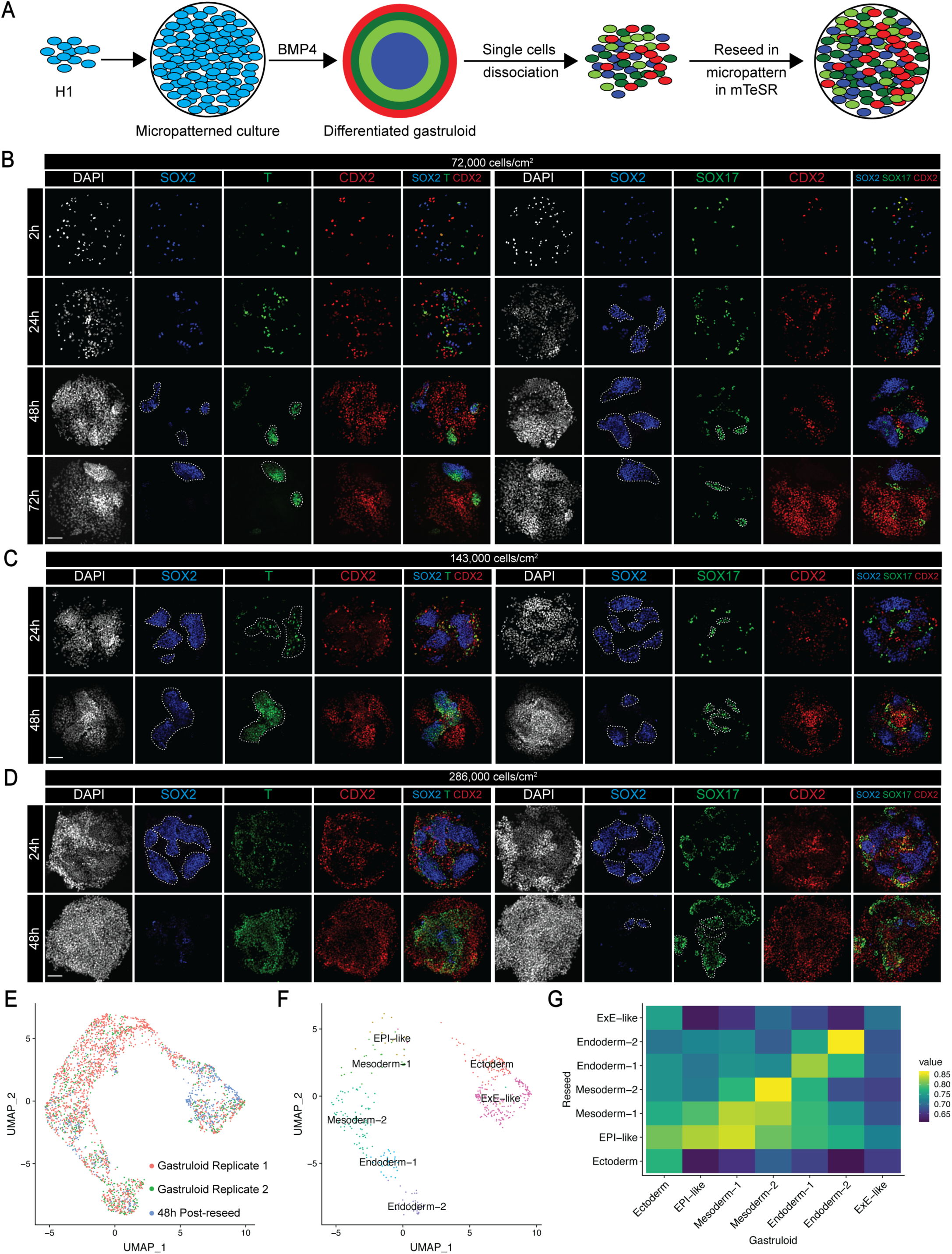
BMP4-differentiated germ layer and ExE-like cells exhibit cell sorting behaviors. (A) Schema of protocol for assaying sorting behaviors of cells from dissociated gastruloids. (B-D) Immunofluorescence images of indicated markers in reseeded gastrloid cells at different timepoints and different cell densities of reseeding. Dashed line indicates cellular aggregates; Scale bar 100 µm. (E) UMAP displaying overlap of two gastruloid replicates and 48h post-reseed cultures. Results of scRNA-seq analysis of ∼36 microcolonies/experiment. (F) UMAP showing clusters of reseeded cultures after integration with gastruloids. (G) Heatmap illustrating gene expression correlation between clusters of reseeded and gastruloid cells.

We next asked whether sorting behaviors depended on cell density by testing gastruloid cells reseeded at low (72,000 cells/cm^2^), medium (143,000 cells/cm^2^), and high (286,000 cells/cm^2^) density (Figure 5B–D). At medium and high densities, we observed aggregation behavior within 24h post-reseeding. The higher the initial reseeded density was, the earlier aggregation behavior was apparent. However, at high density, cells overpopulated the culture discs and aggregation patterns were not as evident as at low or medium densities (Figure 5D). Moreover, immunofluorescence revealed rapid loss of SOX2^+^ by 48h, possibly due to overcrowding of other cell types (Amoyel and Bach, 2014) (Figure S4C). We also observed a slight decrease in SOX2^+^ cells by 48h at medium density, while the number of SOX2^+^ cells remained constant at low density. In contrast, T^+^, SOX17^+^, and CDX2^+^ cells increased in number over the culture period at all densities. Nonetheless, SOX2^+^, T^+^, and SOX17^+^ cells exhibited distinct aggregation patterns at medium density at both 24h and 48h (Figure 5C). Therefore, we further characterized cell aggregation and sorting behaviors in dissociated human gastruloids at medium reseeding density.

We also used scRNA-seq to ask whether the gastruloid cells reseeded in micro-discs for 48h significantly altered their identities from the time of gastruloid dissociation. We performed scRNA-seq on medium-density reseed cultures and analyzed transcriptomes of 479 cells with a mean expression of 4,464 genes (Figure S2A). Using the Seurat package (Butler *et al*., 2018; Stuart *et al*., 2019), we integrated the two gastruloid replicates and reseed cultures and found that all samples overlap closely with each other (Figures 5E, 5F, and S5A), arguing against significant changes in cellular identity between the reseeded and the original gastruloid cultures. Accordingly, based on the identity and expression levels of commonly expressed genes, we found a strong correlation in cluster identities between gastruloid and reseeded microcolonies (Figure 5G). However, we noted changes in proportions of these seven cell types (Figures S5B). Specifically, the proportion of less differentiated cell types such as EPI-like and PS-like Mesoderm-1 decreased in reseeded cultures, whereas those of Mesoderm-2 and Endoderm-2 remained stable. Meanwhile, the proportion of Ectoderm and ExE-like cells increased. Nonetheless, we found that canonical markers used to annotate each gastruloid cluster (Figure 2C) were expressed in respective clusters of reseeded cultures (Figure S5C). Therefore, dissociated gastruloids, upon reseeding, maintained the seven original cell types for 48h, and showed cellular aggregation behavior, a characteristic of cell sorting capability.

### Selective cell sorting of reseeded gastruloid cells

In a typical gastrula, including human, germ layers are arranged so that mesoderm is sandwiched between ectoderm and endoderm, and thus ectoderm lies adjacent to mesoderm but is separated from endoderm (Gilbert, 2010). Similarly, in BMP4-derived gastruloids, the mesoderm ring separates ectoderm from endoderm (Figure 1C) (Warmflash *et al*., 2014). Notably, immunofluorescence images of the 48h reseeded cultures showed that SOX2^+^ ectodermal cells tended to mix with T^+^ mesodermal cells (Figures 6A and S5D–E) but segregated from SOX17^+^ endodermal cells (Figures 6B and S5D). Closer inspection in 3D confocal stacks indicated that SOX2^+^ and T^+^ cells were intermixed and closely associated, with some cells overlapping on top of each other (Figure 6A). Next, we quantified the spatial overlap between pairs of cell types using cosine similarity analysis (Figures 6C and S5F–G). By 24h post-reseeding, the tendency of SOX2^+^ cells to intermingle with T^+^ cells over SOX17^+^ cells was corroborated by significantly higher cosine value between expression domains of SOX2 and T compared to that between SOX2 and SOX17 in medium and high densities reseeded cultures. By 48h, strong association between SOX2^+^ and T^+^ cells was significant at all cell densities. In contrast, the cosine value between expression domains of SOX2 and SOX17 was significantly lower and remained relatively unchanged from 24h to 48h at all cell densities (Figures 6C and S5F–G) despite an increase in SOX17^+^ cells over time (Figures S4C). Overall, higher cosine value between T^+^ mesoderm and SOX2^+^ ectoderm compared to that between SOX17^+^ endoderm and SOX2^+^ ectoderm supports the notion that ectodermal cells more readily associate with mesodermal cells but segregate from endodermal cells.

**Figure 6.**
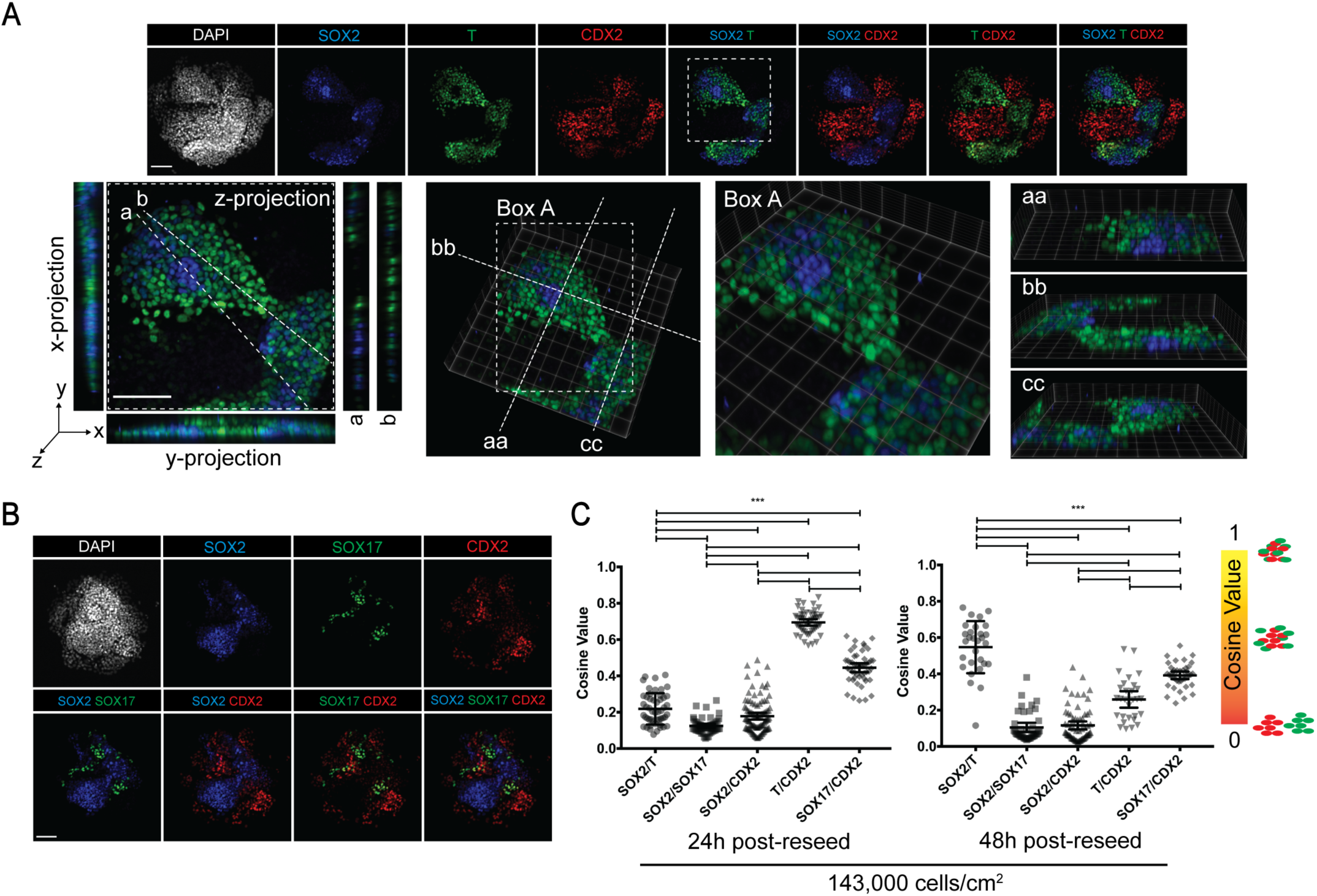
Reseeded gastruloid cells exhibit selective cell sorting behaviors. (A) Immunofluorescence images of indicated markers in 48h post-reseed cultures in (top) 2D and (bottom) 3D. (B) Immunofluorescence images of indicated markers in 48h post-reseed cultures. (C) Cosine similarity analysis of expression domains between pairs of indicated markers; n ≥ 16; *** p < .001; Error bars represent standard deviation. Scale bar is 100 µm.

While cell sorting experiments have been performed in frog and fish gastrulae comprised of the three germ layers (Davis, Phillips and Steinberg, 1997; Ninomiya and Winklbauer, 2008; Klopper *et al*., 2010), it is unclear whether ExE cells sort relative to embryonic cells. As CDX2^+^ ExE-like cells were present in gastruloids, they were included in the reseeding experiments along with germ layer cells, allowing us to test their sorting interactions. Immunofluorescence analyses of 48h post-reseed cultures with anti-CDX2 antibody showed that while CDX2^+^ cells readily intermingled with T^+^ and SOX17^+^ cells, they tended to segregate from SOX2^+^ cells (Figures 6A–B and S5D). This was corroborated by higher cosine value between T^+^ or SOX17^+^ and CDX2^+^ cells compared to that between SOX2^+^ and CDX2^+^ cells across all densities at all time-points (Figures 6C and S5F–G). Together, these experiments reveal the ability of germ layer and ExE-like cells in dissociated gastruloids to sort relative to different cell types to various degrees.

### Reseeded gastruloid cells differentially express cell adhesion molecules

In the cell sorting experiments, we used SOX2, T, SOX17, and CDX2 to identify ectoderm, mesoderm, endoderm, and ExE-like cells, respectively. Towards defining gene expression patterns underlying sorting behaviors, we combined clusters with strong expression of *SOX2* (EPI-like and Ectoderm clusters), *T* (Mesoderm-1 and −2 clusters), *SOX17* (Endoderm-1 and −2 clusters), and *CDX2* (ExE-like cluster), termed reEcto, reMeso, reEndo, and reExE, respectively (Figure S6A–C). We first asked whether differential adhesion could be responsible for aggregation of same cell types by investigating expression of cell adhesion components across cell types. Interestingly, *SOX17^+^* reEndo cluster exhibited the highest expression of classical cadherins (*CDH1*, *CDH2*, *CDH3*), protocadherin PCDH1, and *EPCAM* (> 2-fold over reEcto, reMeso, and reExE clusters). *PCDH10* was also enriched in reEndo and reMeso clusters (> 2-fold over reEcto and reExE clusters). In contrast, *CDH11* was enriched in reEcto, reMeso, and reExE clusters (> 2-fold over reEndo cluster), while *PCDH7* was enriched in reEcto and reExE clusters (> 2-fold over reMeso and reEndo clusters) (Figures S6D and S6E). Future studies will test whether differential expression of these adhesion molecules plays a role in aggregate formation within same cell types of reseeded gastruloid cells.

### Reseeded cells express Ephrin-Eph interacting pairs

Ephrin/Eph signaling has been shown to be responsible for cell sorting or tissue separation in germ layers of *Xenopus* gastrulae (Rohani *et al*., 2011; Fagotto *et al*., 2013). We asked whether the Ephrin/Eph pathway could play a role in the selective cell sorting behaviors of reseeded gastruloid cells. To this end, we used a python package, CellPhoneDB (Efremova *et al*., 2019), to investigate potential interactions between pairs of cell clusters based on the expression of ligands and receptors in respective clusters. CellPhoneDB predicted multiple Ephrin/Eph interacting pairs (Figures S6F and S6G), implying a potential role of the Ephrin/Eph pathway in the selective cell sorting behaviors of reseeded gastruloid cells. In particular, we found strong predicted interactions from reEndo to reEcto (*EPHB6–EFNB3, EPHB6–EFNB2, EPHB3– EFNB3*), as well as those from reEcto to reEndo (*EFNB2–EPHB4/EPHB3/EPHA4*), suggesting that these interactions may play a role in segregation behavior observed between reseeded SOX2^+^ reEcto and SOX17^+^ reEndo cells. Moreover, we found high predicted interactions between *EPHB2* and *EFNB1/EFNB2* from CDX2^+^ reExE to SOX2^+^ reEcto, two cell types which tended to segregate in reseeded cultures. Overall, our experiments revealed that gastruloid cells, when dissociated and reseeded, exhibit cell sorting behavior (Figures 5 and 6), which may be explained by a combination of differential cell adhesion (Figure S6D) and attractive/repulsive interactions mediated by Ephrin/Eph signaling (Figure S6F).

## Discussion

Human gastrulation remains unstudied due to ethical and experimental limitations. However, forward and reverse genetics studies in research animals provide deep insights into germ layer induction, patterning, and morphogenesis that inform hPSC-based experimental platforms to study aspects of human gastrulation (Rossant and Tam, 2016; Simunovic and Brivanlou, 2017; Shahbazi and Zernicka-Goetz, 2018; Taniguchi, Heemskerk and Gumucio, 2019). This work adopts and extends the micropatterned gastruloid system established by Brivanlou and collaborators (Warmflash *et al*., 2014). Our results demonstrate that BMP4 treatment of spatially confined hESCs on ECM micro-discs reproducibly generates radially-arranged cellular rings that resemble the three germ layers and ExE cells (Warmflash *et al*., 2014). Using scRNA-seq and unbiased computational analyses, we reveal a higher cellular complexity of gastruloids that encompasses seven cell types: EPI-like, Ectoderm, Mesoderm-1, −2, Endoderm-1, −2, and ExE-like, and their transcriptomes. Furthermore, we show that upon gastruloid dissociation, the resulting reseeded single cells exhibited cell sorting capabilities *in vitro*, similar to germ layer cells in frog and fish embryos.

Previous studies have used a selection of candidate genes and proteins to define cell types induced by BMP4 in human gastruloids, suggesting they generate four homogenous cell populations (Warmflash *et al*., 2014). Our scRNA-seq data, encompassing 2,475 cells from two independent experiments with multiple gastruloids, uncovered expression of 25,271 transcripts. Comparisons of transcriptomic signatures with mouse (Pijuan-Sala *et al*., 2019) and cynomolgus monkey (Ma *et al*., 2019) gastrulae suggest that the hESC gastruloids resemble E7.25 mouse and 14 dpf cynomolgus monkey gastrulae (Figures 3E–H), and therefore may represent early-to-mid stages of human gastrulation.

Interspecies comparison allowed us to compare gastruloid clusters identified through canonical markers with mouse and monkey cell types based on transcriptomes. Mouse cell types are well-annotated for germ layers and their derivatives across gastrulation stages, enabling us to investigate similarities and subtypes of germ layer cells in gastruloids. Our computational analysis identified an EPI-like cluster resembling mouse epiblast and post-implantation monkey epiblast (Figures 3I and 3J). We also showed the presence of Ectoderm expressing both nonneural and neural ectoderm markers enriched in mouse gastrulae (Pijuan-Sala *et al*., 2019). Moreover, we identified two mesodermal populations, Mesoderm-1 resembling PS and Mesoderm-2 resembling nascent and differentiating mesoderm. We also identified Endoderm-1 which may represent precursors of primitive and definitive endoderm. However, there are known differences between mouse and human in the formation of ExE cells (Shahbazi and Zernicka-Goetz, 2018) and PGC (Tan and Tee, 2019). Accordingly, we did not find similarities between mouse PGC and Endoderm-2, a cluster enriched in *NANOS3*, *TFAP2C*, and other primate PGC and hPGCLC markers. However, cross-species comparison analysis with monkey indicated that Endoderm-2 resembled monkey PGCs (Figure 3J). Future experiments are required to investigate functional competence and mechanisms underlying the specification of hPGCLCs in gastruloids. Additionally, the presumptive TE-layer reported previously in gastruloids (Warmflash *et al*., 2014) expressed both TE and amnion markers (Figure 2C), and matched with both TE and amnion in monkey (Figure 3J). This suggests that the CDX2^+^ outermost gastruloid ring contains precursors of TE-like and amnion-like cells or is a mix of both cell types.

Notably, BMP4 treatment of hESCs within a short period of time (2 to 4 days) results in a diverse cell fates, including hPGCLC and amnion-like cells, whether in 2D micropatterns (Figure 3), disorganized 3D aggregates (Chen *et al*., 2019), or controlled 3D cultures in microfluidics (Zheng *et al*., 2019). It is particularly intriguing that BMP4 co-induces hPGCLC and amnion-like cells in these *in vitro* systems, given that current studies in cynomolgus monkey posit that PGCs are induced in amnion through BMP4, along with WNT3A (Sasaki *et al*., 2016). Highlighting the conserved role of BMP4 in PGC induction in mammals, BMP4 signaling from ExE ectoderm has been reported to instruct epiblast cells toward PGCs in mouse (Lawson *et al*., 1999).

While the monkey dataset used in this analysis contains gastrulating cells, they were classified only as early or late stage (Ma *et al*., 2019). In general, we found similarities between gastruloid germ layers and relevant monkey gastrulating cells (Figure 3J). Interestingly, we found matches of monkey ExE mesenchyme (EXMC) in gastruloid Ectoderm, Mesoderm-2, and Endoderm-1. This requires further investigation but may be explained by the relatively large number of EXMC compared to gastrulating cells in the monkey dataset (Ma *et al*., 2019), confounding the analysis. As more data emerge from nonhuman primates, including well-annotated germ layers, the nature of embryonic and ExE-like cells in human gastruloids can be further investigated. Overall, using canonical makers and cross-species analyses, we defined a rich cellular complexity of gastruloids, containing embryonic and ExE cell types.

An exciting aspect of gastruloid culture is the ability to investigate signaling processes underlying the formation of germ layers and ExE cells in a human system (Siggia and Warmflash, 2018). In that regard, the establishment of a reaction-diffusion system between BMP4 signaling at the edge and its antagonist NOGGIN in the center of gastruloid by 24h is sufficient to produce radial patterning (Etoc *et al*., 2016; Tewary *et al*., 2017). By 44h, we did not observe *NOGGIN* RNA expression, but instead found enrichment for other BMP inhibitors *FST* and *GDF3* in the center (Figure S2J). This suggests a shift in BMP4 antagonists from 24 to 44h to maintain the radial patterning in gastruloids. Alternatively, the BMP4-NOGGIN reaction-diffusion system established at 24h is sufficient for the radial patterning observed at 44h, and expression of *FST* and *GDF3* at 44h indicates a different phase of differentiation.

The BMP, WNT and NODAL signaling cascade is important during mammalian gastrulation and gastruloid formation in micropattern (Chhabra *et al*., 2018; Martyn *et al*., 2018). Specifically, WNT and NODAL pathways synergize to instruct mesoderm differentiation in gastruloids, whereas changes in the duration of BMP signaling alter the position of mesodermal ring (Chhabra *et al*., 2018). Thus, the dynamic interplay of signaling pathways, such as the duration and timing, governs cell fate patterns in gastruloids. Suggesting complex communications across cell types formed in gastruloids, we found multiple predicted interactions between pairs of cell types based on the expression of ligand and receptors using CellPhoneDB (Efremova *et al*., 2019) (Table S2). To further delineate the intricate interplay between various signaling components and their roles in cell fate specification in gastruloids, it will be interesting to combine protein and scRNA-seq analyses over the course of gastruloid differentiation. Such analyses can help infer gene regulatory networks underlying human gastrulation.

During gastrulation, specification of germ layers is followed by their morphogenesis to establish the body plan and organ rudiments. This not only entails cell movements and rearrangements, but also segregation or boundary formation between distinct germ layers or distinct cell types within the same germ layer. Such cell sorting behaviors have been described in frog and fish embryos, where embryonic tissues dissociated into single cells were allowed to interact *in vitro* (Townes and Holtfreter, 1955; Krens and Heisenberg, 2011). However, it is unknown whether mammalian, including human germ layer cells are also capable of cell sorting *in vitro*. We showed that human gastruloids, when dissociated and reseeded as single cells onto the ECM discs, exhibited sorting behaviors (Figures 5 and 6) reminiscent of those reported for amphibians and fish. Specifically, reseeded gastruloid cells were motile and exhibited a random distribution of the four major fates. Over time, immunofluorescence experiments indicated that the reseeded cultures contained cells expressing SOX2, T, SOX17, or CDX2 as the original gastruloids. Likewise, scRNA-seq analyses revealed the seven original cell clusters in the reseeded cultures, although the relative proportion of EPI-like cells was reduced, consistent with their progenitor nature (Figures 5 and S5). Both visual inspection and unbiased cosine similarity of immunofluorescence images indicated that cells showed selective association with and/or avoidance of distinct cell types (Figure 6). In particular, cells expressing a marker formed aggregates with those expressing the same marker. Ectodermal cells associated more readily with mesodermal cells than endodermal cells, similar to spatial arrangement in gastruloids and also to the conserved germ layer arrangement *in vivo*, where ectoderm lies adjacent to mesoderm but separated from endoderm (Gilbert, 2010). Moreover, ExE-like CDX2^+^ cells tended to associate with mesendodermal cells but segregate from ectodermal cells. Therefore, our results indicate that human gastruloid cells have the ability to undergo sorting.

Three main models – differential adhesion, differential cortical tension, and contact inhibition – have been proposed to explain cell sorting (Fagotto, 2015). Our gene expression analysis suggests that cadherins may be involved in the sorting of dissociated gastruloid cells through adhesion mechanism, as indicated by the expression of specific cadherins in different cell types (Figure S6D). Complementary expression of Ephrins and Eph receptors in the segregating cell clusters suggested involvement of contact inhibition cues. Receptor-ligand interaction analysis using CellPhoneDB (Efremova *et al*., 2019) indicated multiple plausible Ephrin-Eph interactions within the same cell type as well as across different cell types (Figure S6F). This is consistent with findings in animal embryonic cells where multiple ligands and receptors are expressed and complex functional interactions across tissues are predicted (Fagotto, Winklbauer and Rohani, 2014; Fagotto, 2015). Future experiments will investigate these possibilities.

Our work provides a rich resource for transcriptomic signature of human-relevant cells at the gastrula stage, and for studies of the cellular and molecular mechanisms of human gastrulation. This resource can be used to identify candidate markers for different cell types in human gastrulae. A combination of protein and transcript analyses provides multi-modal investigation of germ layer formation and interactions between gastruloid cell types. Although the gastruloid system lacks some cell types found in mouse gastrula, such as ExE mesoderm, we found amnion- and PGC-like cells, which are present in monkey but not in mouse. Therefore, our work underscores the utility of this micropatterned hESC culture to investigate aspects of human development (Simunovic and Brivanlou, 2017; Siggia and Warmflash, 2018), but also the significance of studying human and nonhuman primate embryos *in vivo* and *in vitro* cultures to provide a critical reference for interpreting the *in vitro* embryogenesis models.

We anticipate that varying concentration and/or combination of exogenous factors or using hPSCs of different differentiation potential, such as naïve cells, could produce additional cell types relevant to gastrulation. However, it is tempting to speculate that BMP4 treatment of hESCs in simple micropatterned cultures, resulting in the formation of cells resembling amnion, PGC, and the three germ layers, recapitulates species-specific and evolutionarily conserved inductive processes. Future studies involving gene expression profiles over the course of gastruloid formation will allow mapping and comparison of the human developmental timeline and dynamic gene regulatory networks with mouse and nonhuman primates. Such studies will be crucial in delineating the similarities and differences between molecular mechanisms and signaling pathways underlying gastrulation in human and animal models and generating an experimental platform for understanding the pathologies of human embryogenesis.

## Acknowledgements

We appreciate the technical advice on the micropattern culture from Ali Brivanlou, Eric Siggia, and their lab members. We thank Sandra Lam (Washington University School of Medicine in St. Louis) for her guidance and assistance in fabricating PDMS stamps. We thank Sarah Waye and Chuner Guo (Washington University School of Medicine in St. Louis) for constructing parts of the data curation libraries. We would also like to thank Margot Williams (Baylor College of Medicine), Sabine Dietmann, Mariana Beltcheva and Gina Castelvecchi, Angela Bowman, and Thorold Theunissen (Washington University School of Medicine in St. Louis) for helpful comments on the manuscript. This study was funded by Washington University School of Medicine in St. Louis and a grant from the Children’s Discovery Institute to L.S.K., and a Vallee Scholar Award and an Allen Distinguished Investigator Award to S.A.M.

## Author Contributions

Conceptualization, K.T.M., S.A.M., and L.S.K.; Methodology, K.T.M., S.H., Y.F., S.C.G., M.A.A., S.A.M., and L.S.K.; Software, S.H. and M.A.A.; Validation, K.T.M., S.H., and Y.F.; Formal Analysis, K.T.M., L.S.K., and Y.F.; Investigation, K.T.M. and S.A.M.; Data Curation, Y.F. and S.A.M.; Writing – Original Draft, K.T.M. and L.S.K.; Writing – Review & Editing, K.T.M., S.A.M., and L.S.K.

## Materials and Methods

### PDMS stamp fabrication

Standard photolithography methods were used to fabricate the Master, which was used as a template in molding stamps for micro-contactprinting (Alom Ruiz and Chen, 2007; Théry and Piel, 2009). Briefly, the silicon wafer was cleaned by dipping in hydrogen fluoride solution for 1 min and rinsing with water twice. A thin layer of photoresist (SU-8 3000, Micro Chem) was then spin-coated on the wafer at 500 rpm for 10 sec, followed by 3,000 rpm for 30 sec. The wafer with spin-coated photoresist layer was baked at 65°C for 5 min and 95°C for 15 min. The optical mask with desired features was placed in contact with photoresist layer and illuminated with the UV light at 150 – 250 mJ/cm^2^ at 350 – 400 nm. The photoresist layer was then baked at 65°C for 5 min and 95°C for 10 min. Subsequently, the photoresist was developed in developer solution for 5 min. The photoresist Master was then coated with a layer of chlorotrimethylsilane in the vacuum for 30 min. Polydimethylsiloxane (PDMS) and its curing agent (Sylgard 184, Dow Corning) in 10:1 ratio were mixed, degassed, poured over the top of Master, and cured at 60°C overnight, after which the PDMS layer was peeled off to be used as a stamp in micro-contactprinting.

### Microcontact printing

PDMS stamps (approximately 1 cm x 1 cm) were sterilized by washing in ethanol solution and dried in a laminar flow hood. The stamps were treated with O_2_ plasma at 100–150 microns for 2 min. Growth factor reduced Matrigel (Corning) was diluted in DMEM/F-12 (Gibco) at 1:20 dilution and incubated on the stamps to cover the entire surface of the feature side at room temperature for 45 min. Matrigel solution was then aspirated off the stamps, which were air-dried. Using tweezers, Matrigel-coated surface of stamps were brought in contact with glass or plastic substrate for 2 min, ensuring conformal contact between features and substrate. The stamps were then removed and rinsed in ethanol for future uses. Matrigel-printed substrates were incubated with 0.1% Pluronic^TM^ F-68 (Gibco) in DPBS-/- at room temperature for 1 h. In some experiments, we skipped incubation with Pluronic F-68, and did not observe a difference in cell attachment. Finally, the substrates were washed with DPBS-/- for 4 times. Matrigel-printed substrates were stored in DPBS-/- solution at 4°C for up to 2 weeks.

### Cell seeding protocol

Two human embryonic stem cell lines were used in this study: H1 and H9 (WiCell). Both cell lines were routinely cultured on six-well plates coated with growth factor reduced Matrigel in mTeSR media (STEMCELL Technologies) with daily media replacement. The cells were passaged at 80% confluence using Gentle Cell Dissociation Reagent (STEMCELL Technologies) per manufacturer’s protocol. Both cell lines were cultured at 37°C and 5% CO_2_.

For gastruloid differentiation, we adapted the protocol developed by Warmflash *et al*. (Warmflash *et al*., 2014). H1 and H9 cells at 80% confluence were collected by Accutase (Sigma Aldrich) incubation at 37°C and 5% CO_2_ for 8 min, after which equal volume of RPMI medium 1640 (Life Technologies) was added. The cell solution was centrifuged at 300 rcf (relative centrifugal force) for 5min, after which the supernatant was removed. A single cell suspension was then generated with fresh mTeSR. Cells were counted using Countess II Automated Cell Counter (Life Technologies) per manufacturer’s instructions and seeded onto the micropattern at 132,000 – 263,000 cells/cm^2^ in mTeSR with 10 μM Rho-associated kinase inhibitor (ROCKi Y-27632, Millipore Sigma). After 6h, medium containing ROCKi was replaced with mTeSR. The cells were cultured at this stage for 2h, after which the medium was replaced with mTeSR containing 50 ng/mL BMP4 (R&D Systems) for 42–48h.

For cell sorting experiments, gastruloids that have been treated with BMP4 for 44h were collected by Accutase incubation at 37°C and 5% CO_2_ for 10min, after which an equal volume of RPMI medium 1640 was added. A single cell suspension was generated by gently pipetting the cell suspension for 5 times. The cell suspension was centrifuged at 300 rcf for 5min, after which the supernatant was removed. A single cell suspension was then generated with fresh mTeSR. Cells were then counted and seeded onto fresh micropattern at 72,000 cells/cm^2^,143,000 cells/cm^2^, or 286,000 cells/cm^2^ in mTeSR with 10μM ROCKi Y-27632. After 4h, medium containing ROCKi was replaced with mTeSR. mTeSR was replaced daily to wash away unattached cells.

### Immunocytochemistry

Cells were rinsed once with PBS, fixed in 4% paraformaldehyde for 30 min, and rinsed twice with PBS. Blocking solution was made with 0.1% Triton-X and 3% Normal Donkey Serum (Jackson Immunoresearch) in PBS and washing solution was made with 0.1% Tween-20 in PBS. Fixed cells were incubated in blocking solution for 30 min at room temperature before incubation with primary antibodies (see Supplementary Table 1 for primary antibodies used and their dilution) in blocking solution at 4°C overnight. Cells were then washed 3 times in washing solution before being incubated with secondary antibodies (see Table S4 for secondary antibodies used and their dilution) and DAPI at 4 μg/mL (Invitrogen) in blocking solution for 1 h. Finally, cells were washed 3 times in washing solution and stored in PBS or mounted with coverslips using Vectashield Antifade Mounting Medium (Vector Laboratories).

### Microscopy

All confocal images were acquired on Olympus IX81 Inverted Spinning Disk Confocal Microscope with 10X or 20X lenses. Z-stack images of ∼150 µm thick were acquired in four channels corresponding to DAPI, Alexa 488, Alexa 555, and Alexa 647 conjugated antibodies. Each z-stack was projected into a single image for all channels prior to analysis.

### Quantification of fluorescence intensity

Fiji (Schindelin et al., 2012) was used to process and analyze fluorescent microscopic images. We first created masks for each fluorescent image. We then used Concentric Circles plug-in to overlay 20 equally-spaced concentric circles on the image of gastruloid. We measured the average fluorescence intensity along the circumference of each concentric circle. We normalized the fluorescence intensity of each marker of interest to that of DAPI, resulting in normalized average fluorescence intensity value of each marker along 20 different radii of every gastruloid colony. Finally, these values were averaged for multiple colonies per marker and presented.

### Density map-based cell counting and cosine similarity analysis

We estimated the number of cells expressing each marker in a given fluorescence image by first estimating the marker’s corresponding density map, which represents the expected spatial distribution of the number of cells over pixels in the fluorescence image acquired for the marker. Individual 2D fluorescence images were converted into 2D density maps, where the density value of a pixel is the expected number of cells at that pixel in the corresponding image. The density map of a given image *X* ∈ *R^N×N^*, was estimated by use of deeply-supervised fully convolutional neural network (FCNN) models (He *et al*., 2019) that perform an end-to-end mapping from *X* to the desired density map, *Y* ∈ *R^N×N^*. The FCNN model was learned by use of a set of training data, each of which contains an annotated image and the associated ground-truth density maps. Here, each image was manually annotated by identifying the locations of centroids of cells expressing markers of interest. The ground-truth density map of each image was generated based on the annotated centroids by use of methods introduced in previous works (He *et al*., 2019). Specifically, 5 FCNN models, each representing SOX2, CDX2, SOX17, DAPI, and T, were separately trained and developed with corresponding annotated images. In each FCNN model, 7 annotated images with 512×512 pixels were employed as the training images for the FCNN model training, and 3 annotated images were employed to validate select values of parameters in the FCNN models.

Interpreting the density map of an image as the expected spatial distribution of the number of cells over all pixels, the total number of cells in the image is calculated by summing up all the densities on the map:

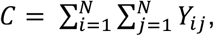

where *Y_ij_* is the estimated density value of the location (*i*, *j*) in the corresponding image. Density map generation or cell counting for CDH2 and GATA3 was performed with FCNN model trained with DAPI and CDX2, respectively.

Representing the density maps corresponding to two cell distributions as *Y* ∈ *R^N×N^* and *Y*^’^ ∈ *R^N×N^*, the spatial overlap between the two cell populations is measured based on the density maps using cosine similarity value (cosine value):

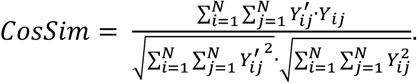

Cosine values ranged from 0 to 1, with 1 representing the highest overlap between expression patterns of two markers, whereas 0 representing no overlap. Intuitively, the cosine value based on density maps is a straightforward way to measure the spatial overlap between two cell populations (cells expressing two markers of interest). According to the definition above, pixels where both of the two cell populations have higher number of cells contribute more to the global overlap; the pixels where only one of the two or neither of them have higher densities may contribute less to the global overlap; when two cell populations have larger amount of cells, but relatively less amount of overlapped pixels, the global overlap is small.

### scRNA-seq and data analysis

Cells were collected by Accutase incubation at 37°C and 5% CO_2_ for 10min. Cell clumps were further broken up into single cells by gently pipetting the cell solution 5 times, after which equal volume of RPMI medium 1640 was added. The cell suspension was centrifuged at 300 rcf for 5 min, after which the supernatant was removed. A single cell suspension was then generated with cold DPBS-/-. Cells were then counted and resuspended at 20,000 cells per 200μL of cold DPBS-/- in centrifuge tubes. For each cell suspension, 800 μL of cold methanol was added dropwise. The final cell suspension was incubated on ice for 15 min and kept at −80°C until use.

To prepare single-cell library, 10x Chromium Single Cell 3’ Library & Gel Bead Kit v2, Chromium Single Cell 3’ Chip Kit v2, and Chromium i7 Multiplex Kit were used according to the manufacturer’s instructions. cDNA libraries were then quantified on the Agilent TapeStation and sequenced on Illumina HiSeq 2500.

The Cell Ranger v.2.1.0 pipeline was used to align reads to the hg19.transgenes genome build and generate a digital gene expression matrix. The Seurat package (v.3.0.2) was used for further data processing and visualization. Default settings were used unless noted otherwise. For gastruloid dataset, cells with a low or high number of genes detected (Replicate 1: ≤ 200 or ≥ 7,500; Replicate 2: ≤ 200 or ≥ 6,000) and cells with high total mitochondrial gene expression (Replicate 1: ≥3%; Replicate 2: ≥2.5%) were removed from further analysis. Consequently, we selected 1,722 cells (out of 1,989) and 753 cells (out of 823) from Replicate 1 and Replicate 2, respectively, that passed quality control for further analysis. For 48h-reseed cultures, cells with ≤ 200 or ≥ 6,000 genes detected and mitochondrial gene expression ≥2% were excluded, resulting in the selection of 479 out of 584 cells for further analysis. After normalizing and scaling the filtered expression matrix to remove unwanted sources of variation driven by mitochondrial gene expression, the number of detected UMIs, and the cell cycle, principal component analysis was performed in Seurat.

The two replicates for gastruloids were integrated based on between-dataset anchors identified by Seurat, and the top 2,000 highly variably expressed genes (HVGs) were included for the integrated expression matrix. Cell clusters were then identified by shared nearest neighbor (SNN) resolution at 0.4 using FindClusters function. Dimensionality reduction was performed using RunUMAP (dim = 1:15). We annotated each cluster based on known markers. Differentially expressed genes (DEG) for each cluster were identified by using FindAllMarkers with threshold settings of 0.25 log fold-change and 25% detection rate. Violin plots were generated with VlnPlot function in Seurat using gene expression from ‘RNA’ assay. Heatmaps were plotted with DoHeatMap function using relative expression (z-score).

Using Seurat, reseed data was integrated with two replicates of gastruloids. We transferred gastruloid cluster annotations to the reseed data using FindTransferAnchors and TransferData functions. AverageExpression function was used to calculate average gene expression for the 2,000 HVGs of RNA data for each gastruloid cluster and each transferred annotation cluster of reseed data. Correlation between reseed and gastruloid clusters was calculated by using Spearman correlation between average gene expression levels of each respective cluster and dataset. The correlations were then visualized using a heatmap. CellPhoneDB (v.2.1.1) (Efremova *et al*., 2019) was used to investigate directionality and cross-talks of interaction between pairs of cell types.

For comparison with published data, each dataset – mouse (Pijuan-Sala *et al*., 2019), monkey (Ma *et al*., 2019), and human amnion embryoid model (Zheng *et al*., 2019) – was normalized, scaled, and merged with human gastruloid object applying Seurat anchor-based integration method. Using default settings, integration of gastruloids with mouse, monkey, and amnion embryoid model was performed using 500, 2,000, and 2,000 anchors, respectively (anchors are defined as pairwise correspondences between gastruloid and mouse, monkey, or amnion embryoid datasets; see (Stuart *et al*., 2019)). Cell type labels from mouse and monkey were transferred to human gastruloid cells by FindTransferAnchors and TransferData functions. For each previously-annotated human gastruloid cluster, we calculated its transferred mouse or monkey cell type composition and identified significant transferred cell types using p-value of randomization test < 0.05. For randomization test for significant transferred cell types of each gastruloid cluster *i* of size *n_i_*, we sampled *n_i_* cells with replacement from all gastruloid cells 1,000 times. For each of the *k^th^* time resampling of size *n_i_*, we calculated *Perc_ijk_*, the percentage of transferred cell types *j* in *n_i_* sampled cells and built a background distribution with *Perc_ij1_*, …, *Perc_ij1000_*. We then computed the p-value for cell type *j* in cluster *i* as:

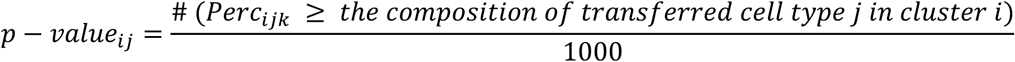

Then for cluster *i*, if *p-value_ij_* is less than or equal to 0.05, we identify transferred cell type *j* as a significant transferred cell type for cluster *i*.

Pairwise Pearson correlation was used to calculate correlation of cell type composition between gastruloid and mouse or monkey embryo at different developmental stages. To calculate correlation between gastruloid clusters and cell types from published datasets, average expression of 2,000 HVGs between gastruloid and each published dataset was calculated. Correlation coefficient between each cell type was calculated using Spearman correlation and plotted in heatmaps.

---

**Figure 1S.**
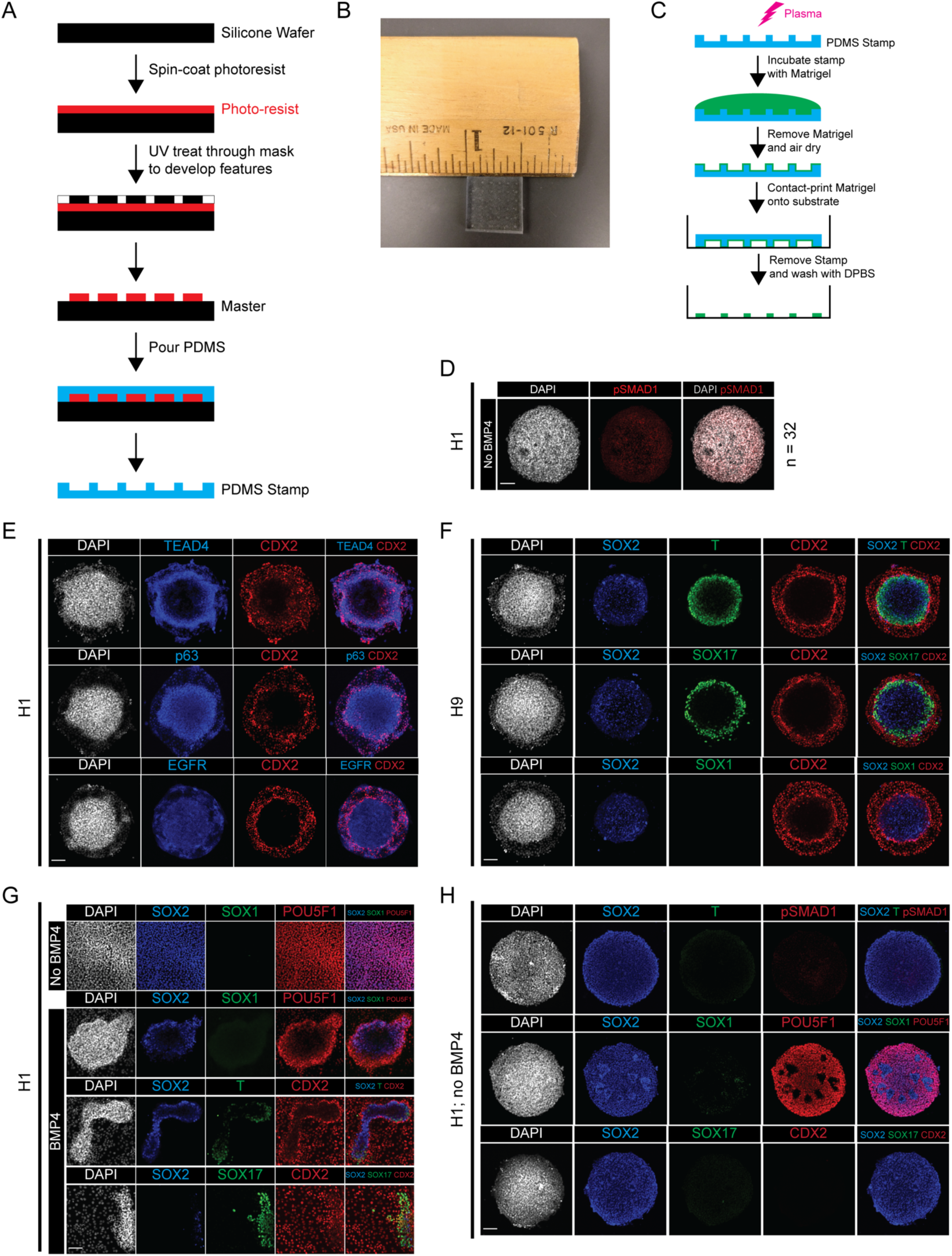
Supplemental information related to Figure 1. (A) Schematic of protocol for fabricating PDMS (Poly-dimethylsiloxane) stamp used in microcontact printing. (B) An image of a typical stamp used in microcontact printing with a ruler and one-inch mark for scale. (C) Schematic of protocol for Matrigel discs microcontact printing using PDMS stamp. (D) Immunofluorescence images of pSMAD1 in micropattern culture of H1 cells without BMP4 treatment (n = 32). (E) Immunofluorescence images of proliferating TE markers in gastruloids differentiated from H1 cells. (F) Immunofluorescence images of indicated markers in gastruloids differentiated from H9 cells. (G) Immunofluorescence images of indicated markers in standard monolayer culture of H1 cells with or without BMP4 treatment. (H) Immunofluorescence images of indicated markers in micropattern culture of H1 cells without BMP4 treatment. Scale bar is 100 µm.

**Figure 2S.**
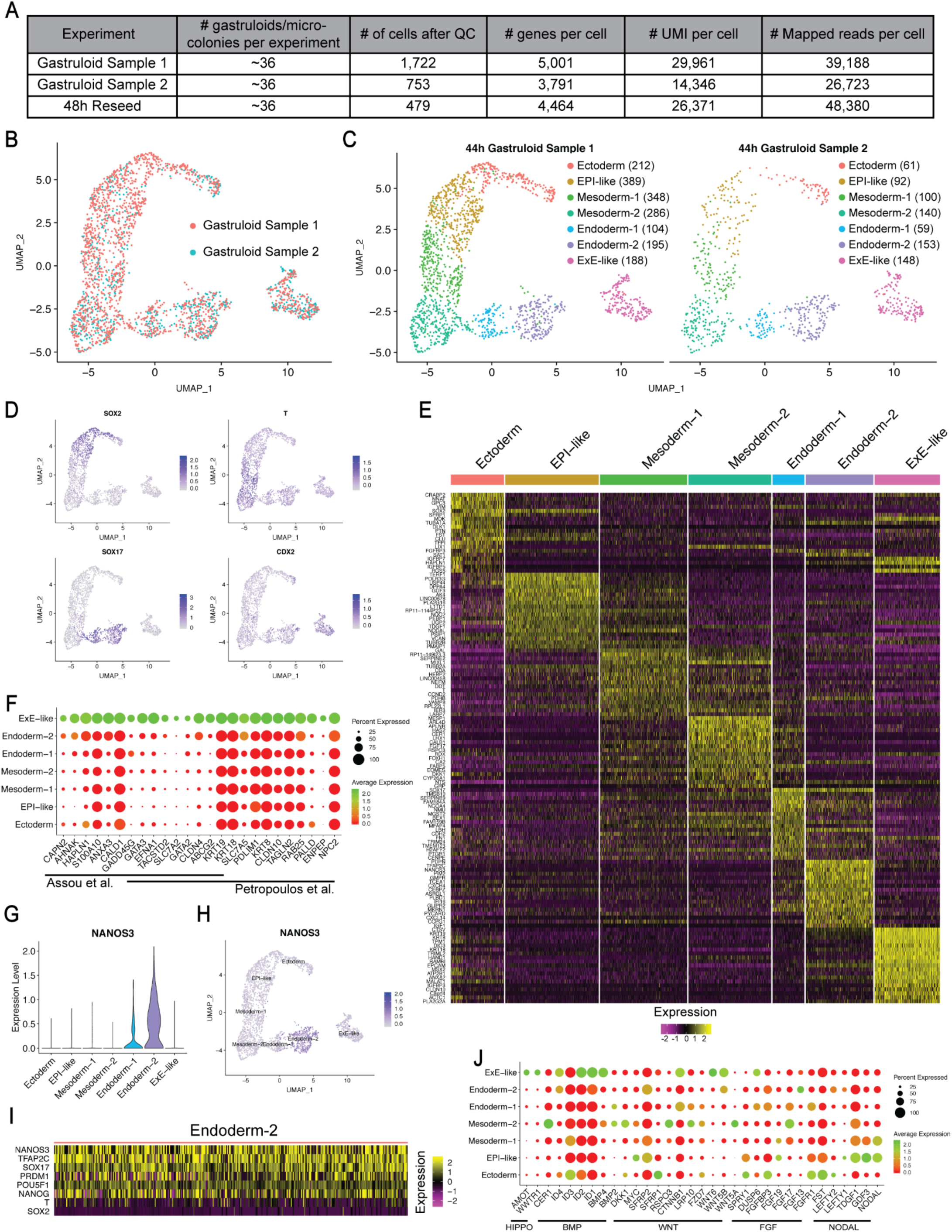
Supplemental information related to Figure 2. (A) Selected parameters of scRNA-seq analysis. (B) UMAP showing overlap of two separate gastruloid scRNA-seq experiments. (C) UMAP display of cluster composition and number of cells in each cluster in the two scRNA-seq experiments. (D) UMAP presenting expression of four makers used in immunofluorescence staining. (E) Heatmap showing top 20 differentially expressed genes in each cluster. (F) Dot plot of TE markers in the gastruloid ExE-like cluster identified in (Petropoulos *et al*., 2016) and (Assou *et al*., 2012). Overlapping bars indicate genes identified in both studies. (G) Violin plot and (H) UMAP illustrating PGC marker *NANOS3* enrichment in Endoderm-2 cluster. (I) Heatmap showing the expression of markers used to identify hPGCLC in Endoderm-2 cluster. (J) Dot plot illustrating expression of HIPPO, BMP, WNT, FGF, and NODAL signaling components.

**Figure 3S.**
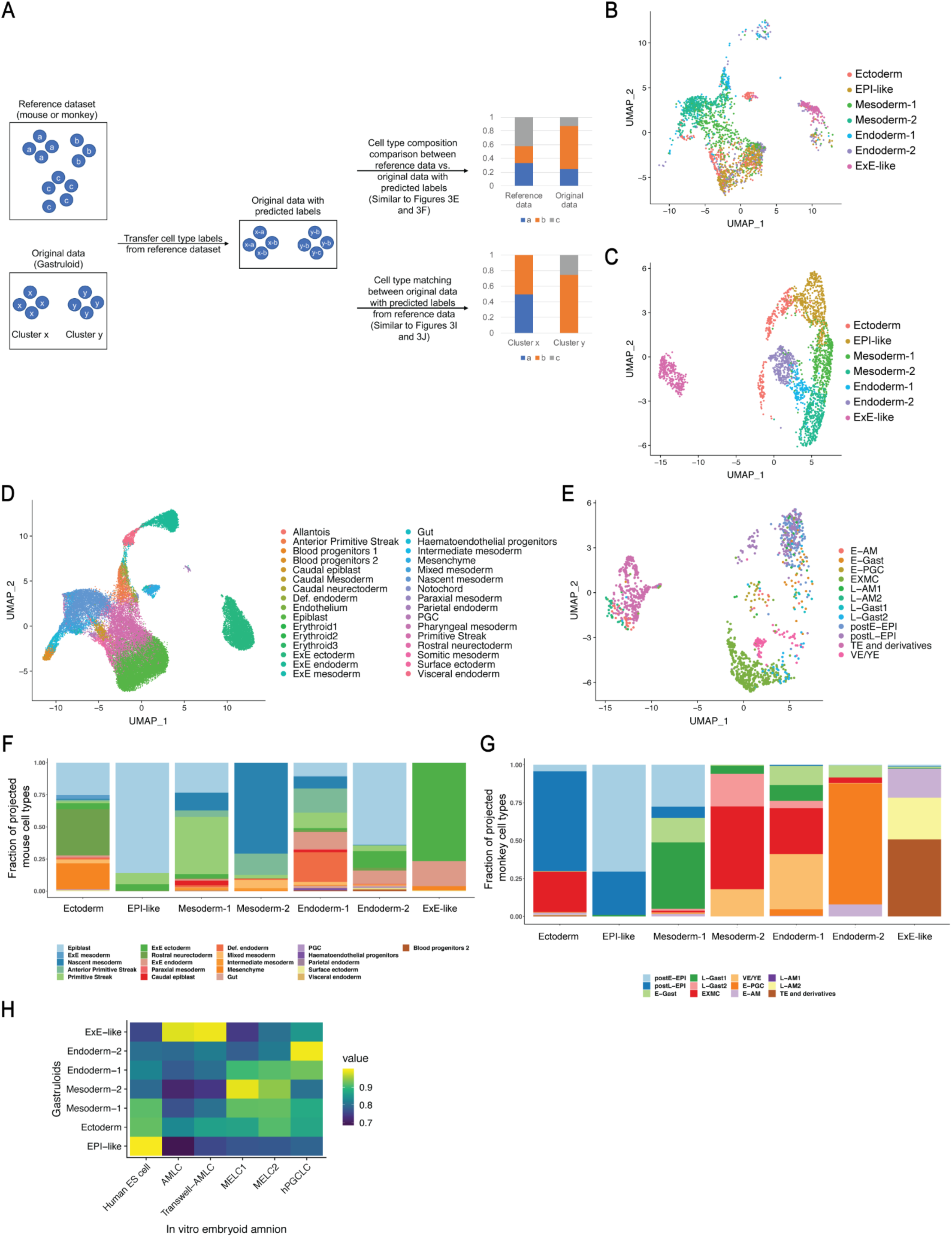
Supplemental information related to Figure 3. (A) Schematic illustrating workflow used for interspecies comparison. (B and C) UMAP displaying gastruloid cells with original cluster annotation after integration with (B) mouse data, and (C) monkey data. (D) UMAP showing mouse gastrulating cells with cell type labels after integration with gastruloid data. (E) UMAP showing monkey post-implantation cells with cell type labels after integration with gastruloid data. (F and G) Bar plots presenting predicted (F) mouse and (G) monkey cell types in each gastruloid cell cluster. Each color bar represents the proportion of a cell type; related to Figure 3I and 3J. (H) Heatmap showing gene expression correlation between the gastruloid cell types and those derived from *in vitro* human embryoid amnion model (Zheng *et al*., 2019). postE/L-EPI, post-implantation early/late epiblast; E/L-Gast, early/late gastrulating cells; EXMC, ExE mesenchyme; VE/YE, visceral/yolk sac endoderm; E-PGC, early PGC; E/L-AM, early/late amnion; AMLC, amnion-like cells; MELC1/2, mesoderm-like cells 1/2; hPGCLC, human primordial germ cell like cells.

**Figure 4S.**
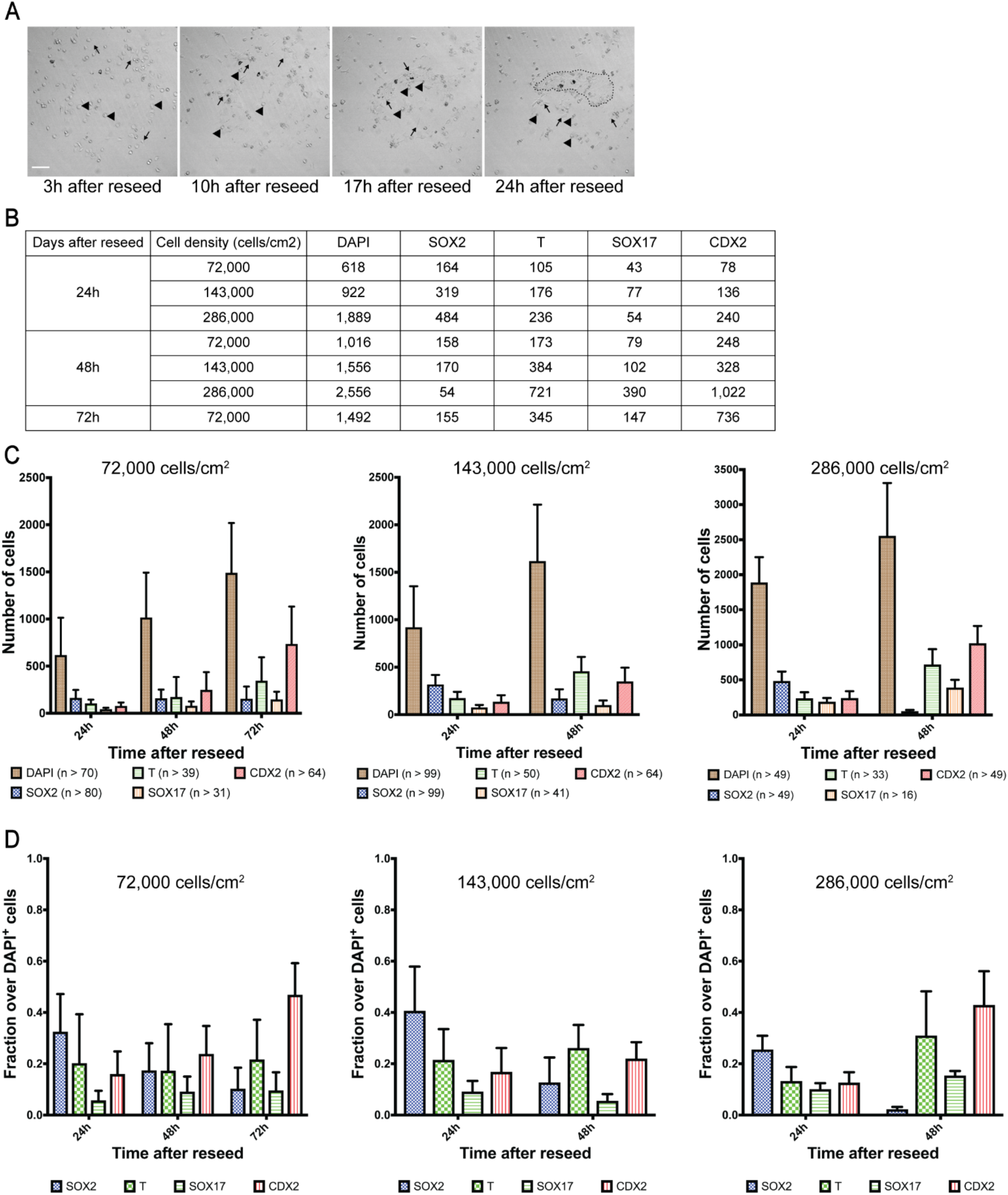
Supplemental information related to Figure 5. (A) Still time-lapse images of dissociated gastruloid cells reseeded in micropattern after 3, 10, 17, and 24h. Arrow indicates motile cells, arrowhead indicates non-motile cells, dotted line indicates cellular aggregate. Scale bar 100 µm. (B and C) Number of dissociated and reseeded gastruloid cells positive for DAPI, SOX2, T, SOX17, and CDX2 at indicated timepoints and cell densities. n ≥ 16; error bars indicate standard deviation. (D) Fraction of dissociated and reseeded gastruloid cells positive for SOX2, T, SOX17, and CDX2 at indicated time points and cell densities.

**Figure 5S.**
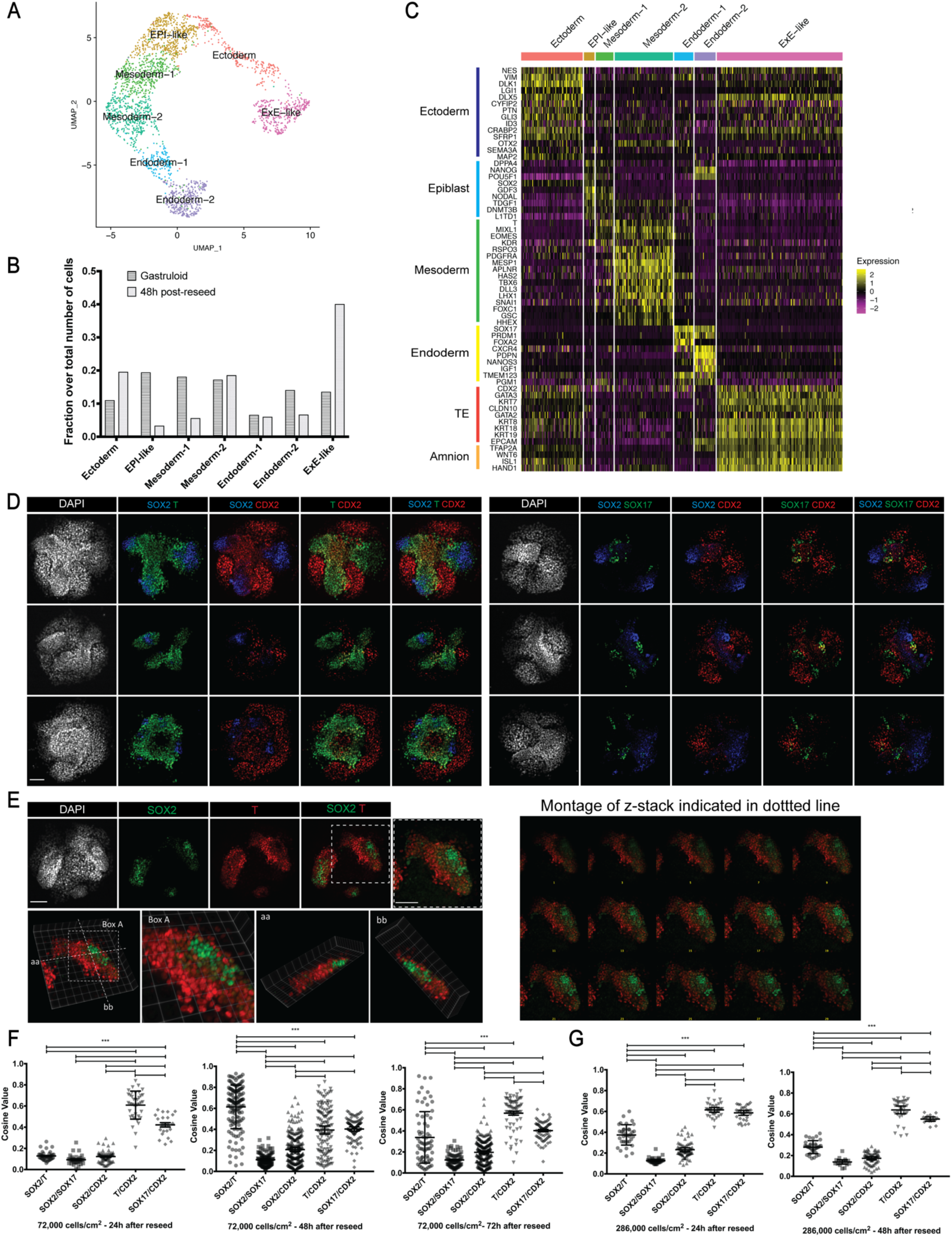
Supplemental information related to Figure 5 and Figure 6. (A) UMAP display of gastruloid cell clusters after integration with reseeded cells. (B) Cellular composition of each cluster in gastruloids and reseeded cell population. (C) Heatmap showing expression of markers used in annotating gastruloid clusters in reseeded cells. (D) Additional replicates of immunofluorescence staining showing indicated markers of dissociated and reseeded gastruloid cells after 48h (note mixing between SOX2^+^ and T^+^ cells, but segregation between SOX2^+^ and SOX17^+^ cells). (E) 2D and 3D-perspective images showing mixing between SOX2^+^ and T^+^ cells in dissociated and reseeded gastruloid cells after 48h. (F and G) Cosine value depicting overlap between expression domains of pairs of indicated markers at indicated timepoints and cell densities. Scale bar is 100 µm.

**Figure 6S.**
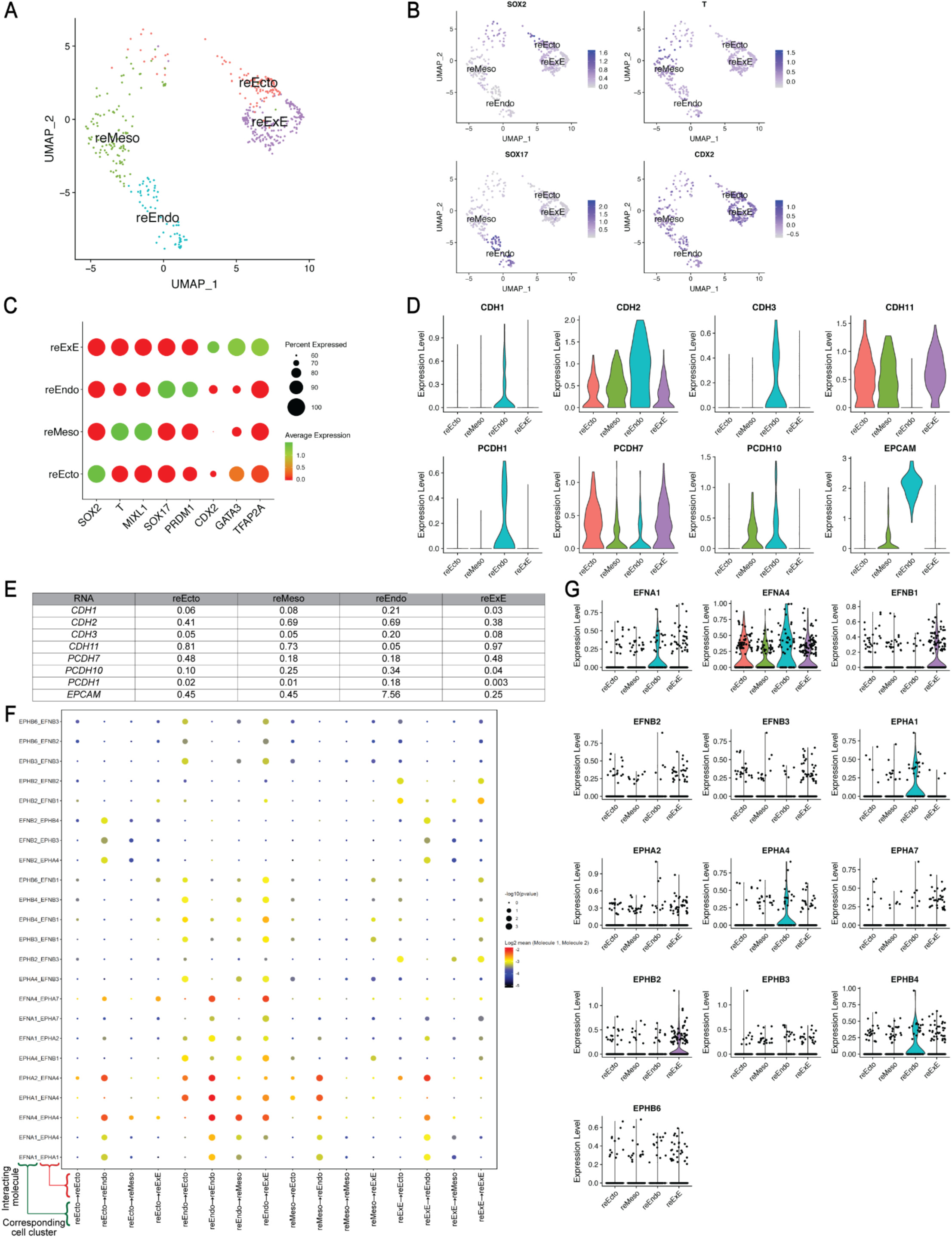
Supplemental information related to Figure 6. (A) UMAP display of reseeded culture from dissociated gastruloids showing combined clusters with strong expression of *SOX2* (EPI-like and Ectoderm clusters into reEcto), *T* (Mesoderm-1 and −2 clusters into reMeso), *SOX17* (Endoderm-1 and −2 clusters into reEndo), and *CDX2* (ExE-like cluster into reExE). (B) UMAP depicting expression of genes encoding the protein markers used in immunofluorescence studies in reseed culture. (C) Dot plot showing marker expression of ectoderm (*SOX2*), mesoderm (*T, MIXL1*), endoderm (*SOX17, PRDM1*), and ExE (*CDX2, GATA3, TFAP2A*) cells. (D) Violin plot depicting expression of indicated cell adhesion markers in the four clusters of reseeded cells. (E) Average expression levels of indicated cell adhesion markers in the four clusters of reseeded cells. (F) Dot plot showing predicted Ephrin-Eph interactions between indicated pairs of cell clusters. Arrows in x-axis labels indicate direction of signaling. In each interacting pair in y-axis, first partner is expressed in the signaling cluster and second partner is expressed in the receiving cluster; *p* value indicates enrichment of predicted interactions; mean indicated average expression of receptor and ligand in interacting cell types. (G) Violin plot illustrating expression of genes encoding Ephrins and Ephs from interaction analysis. Scale bar is 100 µm.

