## Supplemental Table 4 for "High-resolution transcriptional and morphogenetic profiling of cells from micropatterned human embryonic stem cell gastruloid cultures"

| **Primary antibodies** | **Source** | **Dilution** |
| --- | --- | --- |
| anti-SOX2 | Cell Signaling 4900 | 1:200 |
| anti-SOX2 | Cell Signaling 3579 | 1:200 |
| anti-T (Brachyury) | R&D Systems AF2085 | 1:500 (2 μg/mL) |
| anti-SOX17 | R&D Systems AF1924 | 1:1000 (1 μg/mL) |
| anti-CDX2 | Cell Signaling 12306 | 1:100 |
| anti-pSMAD1 | Cell Signaling 9516 | 1:200 |
| anti-ECADHERIN | Cell Signaling 3195 | 1:400 |
| anti-GATA3 | Cell Signaling 5852 | 1:1600 |
| anti-GATA3 | R&D Systems MAB6330 | 1:500 |
| anti-GATA6 | Cell Signaling 5851 | 1:1600 |
| anti-OCT3/4 | Santa Cruz sc-5279 | 1:200 |
| anti-ZO-1 | Invitrogen 33-9100 | 1:500 |
| anti-TEAD4 | Abcam ab58310 | 1:500 |
| anti-CK7 | Cell Signaling 4465 | 1:200 |
| anti-pH3 | Millipore 06-570 | 1:500 |
| **Secondary antibodies** | **Source** | **Dilution** |
| Alexa 488 | ThermoFisher A21202 | 1:500 |
| Alexa 555 | ThermoFisher A21432 | 1:500 |
| Alexa 647 | ThermoFisher A31573 | 1:500 |
